# Structured stabilization in recurrent neural circuits through inhibitory synaptic plasticity

**DOI:** 10.1101/2024.10.12.618014

**Authors:** Dylan Festa, Claudia Cusseddu, Julijana Gjorgjieva

**Affiliations:** School of Life Sciences, Technical University of Munich, Freising, Germany; Max Planck Institute for Brain Research, Frankfurt am Main, Germany; School of Medicine and Health, Institute for Neuroscience, Technical University of Munich, Munich, Germany; Munich Cluster for Systems Neurology (SyNergy), Germany

## Abstract

In cortical circuits, inhibitory interneurons play a dual role: they regulate overall activity levels to prevent runaway excitation, and contribute to diverse computations. While unstructured inhibitory synaptic connections achieve the first role by homeostatically regulating firing rates, computational tasks often require structured excitatory-inhibitory (E/I) connectivity. Here, we consider a broad class of pairwise inhibitory spike-timing dependent plasticity (iSTDP) rules, demonstrating how inhibitory connections can self-organize to both stabilize excitation and generate functionally relevant connectivity motifs—a process we call “structured stabilization”. We show that in both small E/I circuits and large spiking recurrent neural networks the choice of iSTDP rule can lead to either mutually connected E/I pairs, or to lateral inhibition, where an inhibitory neuron connects to an excitatory neuron that does not directly connect back to it. In a one-dimensional ring network with two inhibitory subpopulations following these distinct iSTDP rules, the effective connectivity within the excitatory units self-organizes into a Mexican-hat-like profile, with excitatory influence in the center and inhibitory influence away from the center. This leads to emergent network responses such as contextual modulation effects as in the visual cortex and spatially modular activity characteristic of developmental spontaneous activity. Our theoretical work introduces a family of rules that retains the broad applicability and simplicity of spike-timing-based plasticity, while stabilizing activity and promoting specific connectivity motifs which support emergent network computations.

**Significance Statement:** Neural circuits are faced with the dual challenge of keeping activity stable while modifying synaptic strengths during development and learning. A prominent mechanism to keep excitation in check is inhibitory synaptic plasticity. In recurrent circuit models, we show that plasticity rules based on spike timing at synapses from inhibitory to excitatory neurons can stabilize activity while reinforcing specific connectivity motifs. The shape of these rules dictates the resulting motifs, and determines whether inhibitory neurons form strong reciprocal connections with their excitatory partners or lateral connections onto excitatory neurons that do not connect back. In recurrent networks with local excitatory connectivity, these motifs generate surround suppression as in the visual cortex, and long-range spatial correlations characteristic of developmental spontaneous activity.

## 1 Introduction

In cortical circuits, GABA-ergic interneurons constitute approximately 10% to 20% of the total neural population, and can be further divided into subtypes that exhibit distinct characteristics in connectivity, plasticity and tuning properties (Jiang et al. 2015; Tremblay et al. 2016). Fully understanding how these diverse interneuron subtypes influence neural circuit function remains a significant challenge. A key open question is how activity-dependent plasticity of inhibitory synapses in recurrent circuits sculpts this diversity into specific connectivity motifs while maintaining stable network activity.

The dynamic balance between excitation and inhibition is crucial for generating the irregular, Poisson-like firing patterns commonly observed *in vivo* in cortical circuits (Van Vreeswijk and Sompolinsky 1998; Brunel 2000). This balance enhances the circuit’s sensitivity to input, facilitating rapid information processing (Tsodyks and Sejnowski 1995). Furthermore, it supports essential computations, such as receptive field emergence (Wallenstein and Hasselmo 1997; Froemke et al. 2007; Dorrn et al. 2010; Kleberg et al. 2014; D’amour and Froemke 2015; Rubin et al. 2017), neural assembly formation (Sadeh and Clopath 2021), and the storage of precise temporal firing sequences (Maes et al. 2021; Riquelme et al. 2023; Gong and Brunel 2024; Shao et al. 2024).

Excitatory-inhibitory (E/I) balance can be achieved without any specific inhibitory connectivity structures. Often circuit models consider inhibition simply as a “blanket” that regulates excitation, and incorporate it into a global normalization term or a correction to the synaptic weights between excitatory neurons (Jin 2002; Vogels et al. 2011; Ravid Tannenbaum and Burak 2016; Montangie et al. 2020; Cusseddu et al. 2026). While this simplified approach captures fundamental properties of neural dynamics, it overlooks the possible advantages of more specific E/I connectivity patterns. Incorporating structured inhibitory connections can further refine computations and circuit properties such as E/I co-tuning (Clopath et al. 2016; Mackwood et al. 2021; Miehl and Gjorgjieva 2022; Eckmann et al. 2024), memory storage (Mongillo et al. 2018; Bergoin et al. 2023; Miehl et al. 2023; Agnes and Vogels 2024; Meissner-Bernard, Jenkins, et al. 2025; Meissner-Bernard, Zenke, et al. 2025), predictive coding (Schulz et al. 2021), and other aspects of neural dynamics (Mongillo et al. 2018; Dahmen et al. 2020; Sadeh and Clopath 2020; Hafizi et al. 2022).

Experimental findings demonstrate that inhibitory neurons form non-random connectivity patterns that cannot be explained solely by cell distance or neuron morphology. Additional factors, including developmental factors such as spontaneous activity, interneuron subtype, synaptic dynamics and targeting preference significantly shape this connectivity structure (Peng et al. 2021; Udvary et al. 2022; Bollmann et al. 2023; Reimann et al. 2023; Wu et al. 2023; Kuan et al. 2024; Lakhera et al. 2024; Znamenskiy et al. 2024; Harth et al. 2025). In the neonatal mouse cortex, even before the onset of external sensory inputs, inhibitory hub neurons regulate circuit development (Bollmann et al. 2023). Moreover, theoretical studies suggest that spatial organization with a widespread Mexican-hat-like effective connectivity, consisting of local excitation and longer-range inhibition, can give rise to modular activity patterns with long-range spatial correlations propagating throughout the neonatal ferret cortex (Smith et al. 2018). Such correlated activity is thought to serve as a scaffold for the emergence of specific stimulus feature tuning in the early stages of development (Mulholland et al. 2021; Lakhera et al. 2024; Trägenap et al. 2025).

Despite these advances, the mechanisms and principles that enable the formation and maintenance of inhibitory connectivity patterns, especially in the context of excitatory connectivity, remain elusive. Genetic mechanisms play a fundamental role, particularly in early developmental stages (Duan et al. 2020; Hanganu-Opatz et al. 2021; Wu et al. 2023). Equally important is synaptic plasticity driven by neural activity, which refines inhibitory connectivity through a dynamic, activity-dependent process (Sreenivasan et al. 2022). This mechanism enables a closed-loop regulation of neural dynamics, allowing for precise tuning and adaptability in the face of change.

While the majority of experimental and theoretical studies investigate excitatory plasticity (e.g. Kempter et al. 1999; Song et al. 2000; Morrison et al. 2007; Shulz and Feldman 2013), far fewer examine inhibitory plasticity. Although some theoretical work has considered the impact of different forms of iSTDP in feedforward circuit architectures with a single postsynaptic neuron (Kleberg et al. 2014; Luz and Shamir 2014), there is still no general understanding of how pairwise iSTDP might influence the formation of specific connectivity motifs at the level of recurrently connected circuits. Theoretical work has proposed an iSTDP rule that guarantees the homeostatic regulation of neuronal rates (Vogels et al. 2011), becoming the “canonical” choice for plastic circuits. However, this rule tends to over-regulate rates in a homeostatic manner, producing narrow rate distributions in the excitatory population, which contradicts experimental findings Buzsáki and Mizuseki 2014. In other work, inhibitory rules are specifically fit to experimental measurements (Lagzi et al. 2021; Bergoin et al. 2025), derived normatively from computation objectives (Mackwood et al. 2021), or are found through numerical optimization on a large parameter space (Confavreux et al. 2024). Here we ask how different spike timing-based rules at inhibitory synapses shape which excitatory and inhibitory neurons become strongly connected in recurrent circuits, and how the resulting excitatory-inhibitory motifs stabilize network firing rate, while supporting surround modulation and the generation of spatially correlated spontaneous activity.

To address these questions, rather than studying single rules, here we propose an integrated approach by analyzing a broad family of pairwise iSTDP rules and study their impact in small analytically tractable E/I circuits as well as in large-scale recurrent neuronal networks (RNNs) of spiking neurons. We identify mechanisms that stabilize excitatory activity while shaping circuit motifs, and show how, depending on the profile of the rule considered, inhibitory connectivity adjusts to the existing excitatory connectivity in distinct ways.

We first consider a recurrently connected two-neuron circuit of one excitatory and one inhibitory neuron, and derive how the choice of the iSTDP rule determines which features of neural activity drive the plasticity of inhibitory connections. At one extreme, rules dominated by firing rates are weakly influenced by existing connectivity structure in the two-neuron circuit, aiming primarily at the homeostatic regulation of firing rates, and thus promoting “blanket” inhibition. At the opposite extreme, we find covariance-dominated rules, strongly influenced by pairwise spike correlations that, in the absence of external inputs, arise from the underlying circuit connectivity. These rules stabilize neural activity, while promoting specific connectivity motifs, such as a mutually connected E/I pair or lateral inhibition, depending on the iSTDP features. We call the emergence of these specific connectivity motifs “structured stabilization”. This structured stabilization also generalizes to large-scale recurrently connected circuits where the same type of E/I connectivity motifs were observed under the different iSTDP rules. To further explore the functional consequences of covariance-dominated rules, we implement an RNN whose excitatory neurons follow a one-dimensional ring topology, and whose inhibitory neurons follow two distinct iSTDP rules. Here, inhibitory connectivity self-organizes into an effective Mexican-hat profile, generating long-range modular correlations of spontaneous activity. Finally, we demonstrate that the circuit implements contextual modulation, including suppression of responses to matching stimuli and amplification for mismatched stimuli. In sum, our study investigates how distinct inhibitory plasticity rules shape E/I connectivity motifs, from mutual to lateral inhibition, giving rise to network response properties such as surround modulation and long-range activity correlations observed in the adult and developing visual cortex.

## 2 Results

### 2.1 Inhibitory plasticity shapes specific E/I circuit motifs

To understand how inhibitory plasticity influences circuit connectivity patterns, we first developed an analytical framework to link the shape of iSTDP to the pattern of learned inhibitory connections in a minimal recurrent E/I circuit. Rather than choosing a single rule, we considered a family of iSTDP rules that depend exclusively on spike times applied to a synapse from an inhibitory presynaptic to an excitatory postsynaptic neuron (inh-to-exc) (Kempter et al. 1999; Luz and Shamir 2014; Akil et al. 2021). Each rule in this family contains two fixed components triggered by each pre- and post-synaptic firing events, denoted as *α*_pre_ and *α*_post_, and a kernel function, *L*(*τ*) triggered by the co-occurrence of pre-post (*τ* > 0) or post-pre (*τ* < 0) spiking events (see Methods, Section 4.1). When the spiking activity is modeled as a stochastic process, the mean-field equation for the evolution of the average synaptic weight can be decomposed as follows (Kempter et al. 1999):

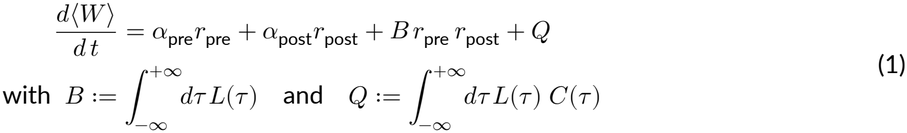

where *r*_pre_ and *r*_post_ are the pre- and post-synaptic firing rates, *B* is the net area of the iSTDP kernel *L*(*τ*), and *Q* captures the effect of the iSTDP kernel on the pre-post pairwise interactions, conveyed by the covariance density curve *C*(*τ*) (see Supporting Information, Section S1.1, or Kempter et al. 1999; Morrison et al. 2008; Luz and Shamir 2014; Lagzi and Fairhall 2024). Eq. 1 shows three types of contributions: terms that depend solely on pre- and post-synaptic firing rates (α_pre_ *r*_pre_ and *α*_post_ *r*_post_), a term that depends on the rates and on the kernel area (*B r*_pre_ *r*_post_), and a term that depends on spike covariance (*Q*). Based on the magnitude of the kernel area *B*, we can distinguish two regimes within this family of rules: a “rate-dominated” regime and a “covariance-dominated” regime.

The first regime, which we refer to as “rate-dominated”, corresponds to rules in the family where *B* is high. In this subset of rules, the contribution of the *B r*_pre_ *r*_post_ term can easily be an order of magnitude above *Q*, even in the presence of strong pre-post correlations. Therefore, when *Q* ≪ *B r*_pre_ *r*_post_, the plasticity dynamics of Eq. 1 are strongly dominated by the firing rates and only weakly driven by the spiking covariance (see Methods, Section 4.3). A prominent example of this is the previously established and well-studied rate-homeostatic iSTDP rule (Vogels et al. 2011). With *α*_post_ = 0 and kernel integral *B* > 0, this rule drives the inhibitory weight to adapt continuously so that the postsynaptic rate converges to the target value 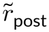, corresponding to a stable fixed point of Eq. 1, namely: 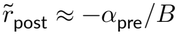.

The second regime, which we refer to as “covariance-dominated”, corresponds to rules in the family for which the iSTDP kernel has zero or near-zero net area (*B* ≈ 0). These rules amplify the contribution of the second-order correlations between pre- and postsynaptic neurons in the *Q* term. In choosing specific pairwise interaction kernels, we considered recent experimental findings that reported a symmetric iSTDP rule for parvalbumin-expressing (PV) interneurons, as well as an antisymmetric iSTDP rule for somatostatin-expressing (SST) interneurons, both projecting onto excitatory pyramidal neurons in the orbitofrontal cortex (Lagzi et al. 2021). Consequently, we focused on symmetric kernels with strong short-range potentiation and weak long-range depression, or antisymmetric kernels in which pre-post and post-pre interactions balance each other with opposite signs. Both kernels can be defined through combinations of exponential shapes, see Eq. 5 for a formal definition.

Analyzing the full dynamics of Eq. 1 in a recurrently connected circuit is challenging because the covariance *C*(*τ*) generally depends on the synaptic weights themselves. We therefore considered a minimal two-neuron E/I recurrent circuit (Fig. 1A) with fixed exc-to-inh connection (*w*_exc_) but plastic inh-to-exc weight (*w*_inh_). We modeled the neurons as time-varying Poisson processes, where each incoming spike changes the instantaneous spiking probability through an excitatory or inhibitory post-synaptic potential (EPSP/IPSP) kernel (internal traces shown in Fig. S1, see also Methods Section 4.4). The inhibitory neuron’s firing rate is kept constant by an external homeostatic mechanism. The excitatory neuron receives an excitatory fixed external input in addition to the inhibitory input. Using this simplified model, we derived analytically a closed-form expression for the fixed-point of the iSTDP dynamics (Methods, Eq. 11; Supporting Information). The condition of fixed *r*_inh_ is not required for a stable solution. Allowing *r*_inh_ to vary still leads to an analytic expression for the fixed point of the iSTDP dynamics, but the resulting closed-form involves the root of a quadratic equation and is therefore less intuitive (see Supporting Information, S1.3, Fig. S3).

**Figure 1:**
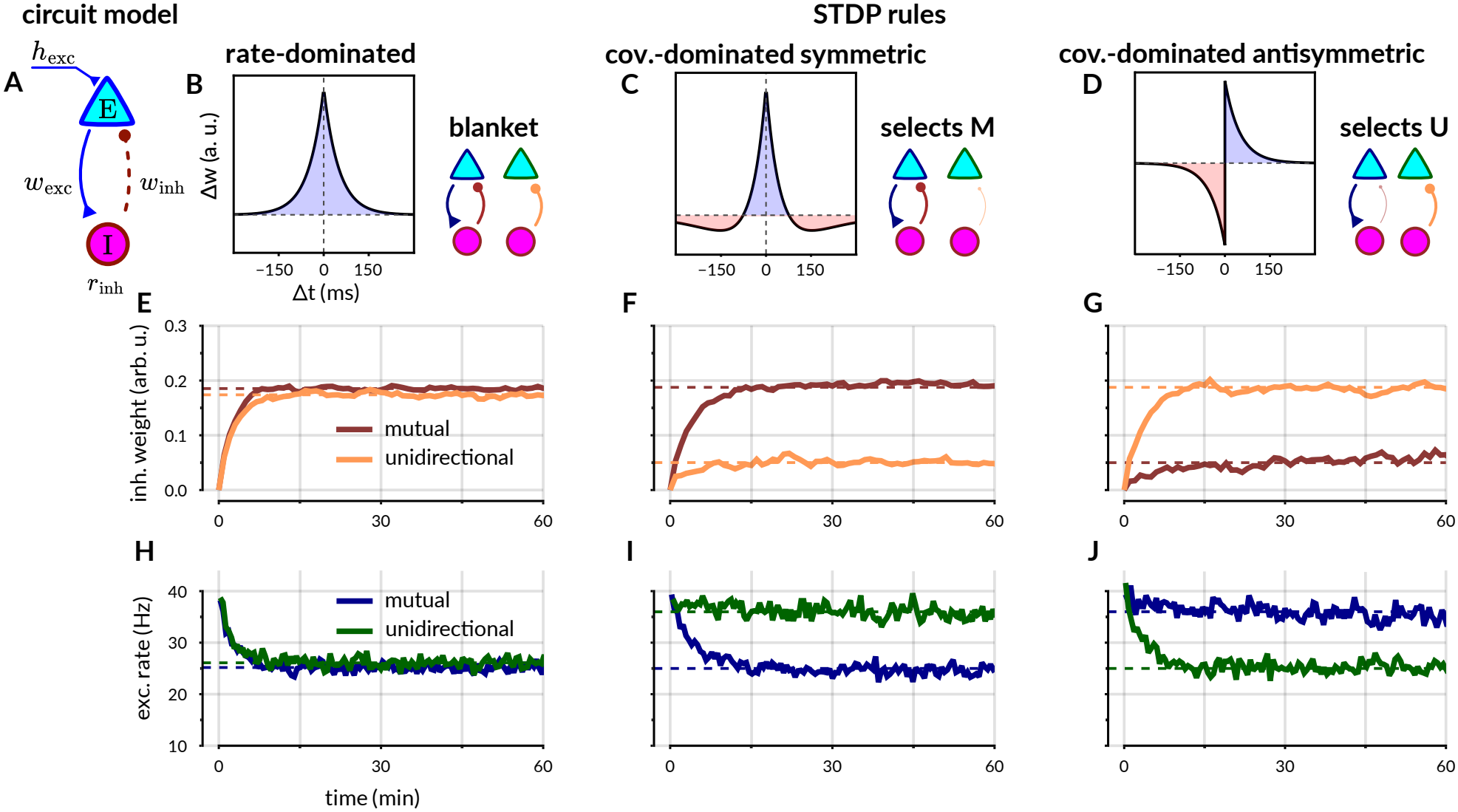
Effects of plasticity on E/I connectivity motifs in a two-neuron circuit. **A.**Schematics of the two-neuron circuit model. Two time-varying coupled Poisson units, one excitatory and one inhibitory, are connected by a fixed excitatory weight *w*_exc_ and a plastic inhibitory weight *w*_inh_. The instantaneous rates are computed as the convolution between each incoming spike and an exponential synaptic kernel that either increases (exc) or decreases (inh) the instantaneous firing probability. The input to the excitatory neuron *h*_exc_ is fixed, whereas the inhibitory input changes dynamically, keeping the inhibitory neuron at a fixed rate *r*_inh_. Only the inhibitory synapse (dashed line) is plastic. The circuit is simulated in two configurations: unidirectional (U), with *w*_exc_ = 0 and mutual (M), with *w*_exc_ = 0.5. **B**,**C**,**D**. Pairwise interaction kernels of the three iSTDP rules tested (Eq. 1). Plots show only the pairwise interaction component. For rate-dependent components *α*_pre_, *α*_post_ and full numerical parameters see Table 1. **E**,**F**,**G**. Change in inhibitory synaptic weight over time for the M (brown) and U (light orange) circuit configurations, under the rules above each column. Dashed lines are analytic predictions for the fixed point (Methods, Eq. 11). **H**,**I**,**J**. Change in excitatory firing rate over time for the M (dark blue) and U (dark green) configurations, under the rules above. Dashed horizontal lines are the analytic predictions for the fixed-point (Methods, Eq. 11).

With this analytical framework in hand, we investigated how different iSTDP rules affect the resulting inhibitory synaptic strength and the excitatory postsynaptic activity. We considered two configurations of our E/I circuit: (1) a unidirectional (U) configuration, where the excitatory-to-inhibitory weight *w*_exc_ is set to zero, and (2) a mutual (M) configuration, featuring a strong excitatory-to-inhibitory connection. We studied the three distinct iSTDP rules in both configurations (whose pairwise interaction kernels *W* (Δ*t*), excluding the rate-dependent terms *α*_pre_ and *α*_post_, are shown in Fig. 1B–D): a rate-dominated rule with a symmetric interaction kernel (Fig. 1B), a covariance-dominated rule also with a symmetric kernel (Fig. 1C), and a covariance-dominated rule with an antisymmetric kernel (Fig. 1D). As expected, the rate-dominated rule shows minimal sensitivity to the presence of the mutual connection. In both U and M configurations, the inhibitory synapse adjusts the excitatory firing rate toward the target rate dictated by the rate-dependent components, approximately given by 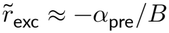 (Fig. 1E). Minor deviations arise from the non-zero contribution of the *Q* term.

**Table 1:**
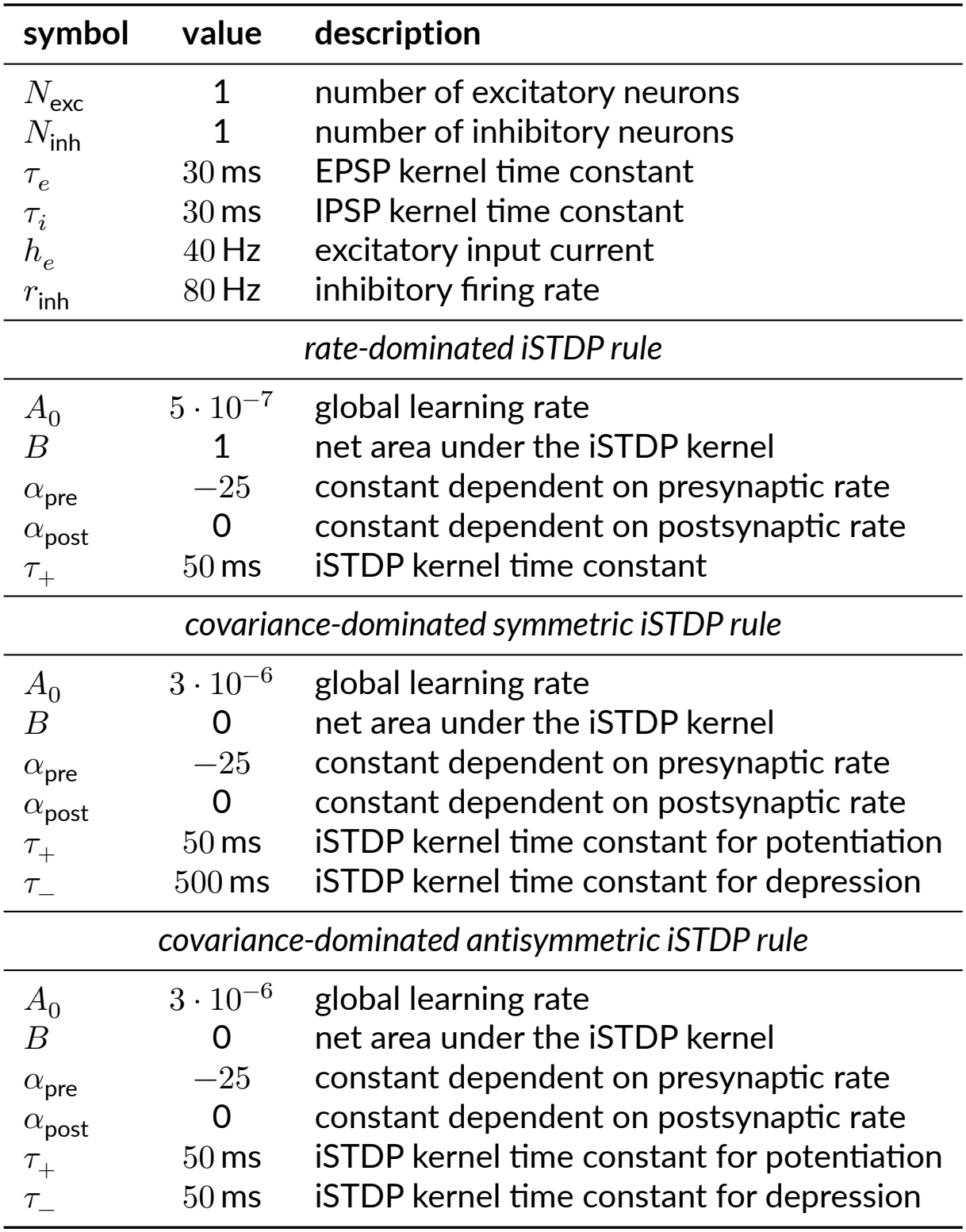
Numerical parameters used for Fig. 1. The parameters in Fig. 2 match those of Fig. 1 unless otherwise indicated in the figure caption.

| symbol | value | description |
|---|---|---|
| N<sub>exc</sub> | 1 | number of excitatory neurons |
| N<sub>inh</sub> | 1 | number of inhibitory neurons |
| τ<sub>e</sub> | 30 ms | EPSP kernel time constant |
| τ<sub>i</sub> | 30 ms | IPSP kernel time constant |
| h<sub>e</sub> | 40 Hz | excitatory input current |
| r<sub>inh</sub> | 80 Hz | inhibitory firing rate |
| *rate-dominated iSTDP rule* |  |  |
| A<sub>0</sub> | 5 · 10<sup>−7</sup> | global learning rate |
| B | 1 | net area under the iSTDP kernel |
| α<sub>pre</sub> | −25 | constant dependent on presynaptic rate |
| α<sub>post</sub> | 0 | constant dependent on postsynaptic rate |
| τ<sub>+</sub> | 50 ms | iSTDP kernel time constant |
| *covariance-dominated symmetric iSTDP rule* |  |  |
| A<sub>0</sub> | 3 · 10<sup>−6</sup> | global learning rate |
| B | 0 | net area under the iSTDP kernel |
| α<sub>pre</sub> | −25 | constant dependent on presynaptic rate |
| α<sub>post</sub> | 0 | constant dependent on postsynaptic rate |
| τ<sub>+</sub> | 50 ms | iSTDP kernel time constant for potentiation |
| τ<sub>−</sub> | 500 ms | iSTDP kernel time constant for depression |
| *covariance-dominated antisymmetric iSTDP rule* |  |  |
| A<sub>0</sub> | 3 · 10<sup>−6</sup> | global learning rate |
| B | 0 | net area under the iSTDP kernel |
| α<sub>pre</sub> | −25 | constant dependent on presynaptic rate |
| α<sub>post</sub> | 0 | constant dependent on postsynaptic rate |
| τ<sub>+</sub> | 50 ms | iSTDP kernel time constant for potentiation |
| τ<sub>−</sub> | 50 ms | iSTDP kernel time constant for depression |

In contrast, the covariance-dominated rules are sensitive to both the circuit’s connectivity structure and the shape of iSTDP kernel. With the symmetric kernel, the final inhibitory weight becomes significantly stronger in the M configuration compared to the U configuration (Fig. 1F, compare brown and light orange). Conversely, with the antisymmetric kernel, the inhibitory weight is lower in the M configuration (Fig. 1G). Such specificity enables these rules not only to regulate excitatory rates, but also to shape specific E/I connectivity motifs—an effect we termed “structured stabilization”. This property emerges from the well-known interaction between the iSTDP kernel and second-order spiking statistics. While this has been previously studied for excitatory plasticity (Kempter et al. 1999; Song et al. 2000; Babadi and Abbott 2013; Shulz and Feldman 2013), here we extend it to inhibitory plasticity in a minimally recurrent two-neuron E/I circuit and hence consider how recurrent inhibition regulates excitation. Specifically, the strong excitatory-to-inhibitory connection in the M configuration generates positively-correlated post-pre events. For the symmetric iSTDP, these events occur within the positive region of the kernel, leading to synaptic potentiation. Conversely, with the antisymmetric iSTDP rule, the same positively-correlated events interact with the negative region of the kernel, resulting in synaptic depression. Additionally, the inhibitory-to-excitatory weight generates a negative covariance density, localized instead in the pre-post region of the curve. As both rules are positive here, this introduces a negative feedback in both cases, limiting the value that *w*_inh_ can reach before plasticity switches into depression. Finally, adjusting the fixed components of the iSTDP rule, *α*_pre_ and *α*_post_, is also crucial for reaching the desired excitatory firing rate (set to 25Hz) in the preferred circuit configuration.

Previous work has shown that rate-homeostatic iSTDP can also generate structured inhibitory connectivity through its sensitivity to spike correlations (Vogels et al. 2011; Lagzi and Fairhall 2024). To investigate this effect within our framework, we extended the two-neuron circuit to include a second inhibitory neuron (Supporting Information, Section S2 and Fig. S4). When calibrated to achieve the same target firing rate and timescale of rate stabilization, both rules converge to the same inhibitory weights, reinforcing the mutually connected E/I motif. However, the timescale of structure formation differs considerably. While the rate-dominated rule initially promotes blanket inhibition and differentiates the inhibitory weights on a slower timescale, the covariance-dominated rule achieves rate stabilization and structure formation on similar timescales.

### 2.2 The shape of inhibitory STDP affects inhibitory connectivity

Having established that covariance-dominated iSTDP rules can stabilize excitatory activity while organizing inhibitory weights to depend on the circuit’s excitatory structure, we next investigated how these outcomes depend on the shape of the iSTDP kernel. We considered four kernel shapes: symmetric, antisymmetric, and their “inverted” counterparts, obtained by reversing the sign of the kernel (Fig. 2A). We analyzed the behavior of these kernels in the reduced two-neuron E/I circuit. To explore the parameter space, we varied the rate-dependent parameters *B* and *α*_pre_, while keeping *α*_post_ = 0. With this parametrization, the covariance-dominated regime corresponds to values of *B* and *α*_pre_ close to zero.

**Figure 2:**
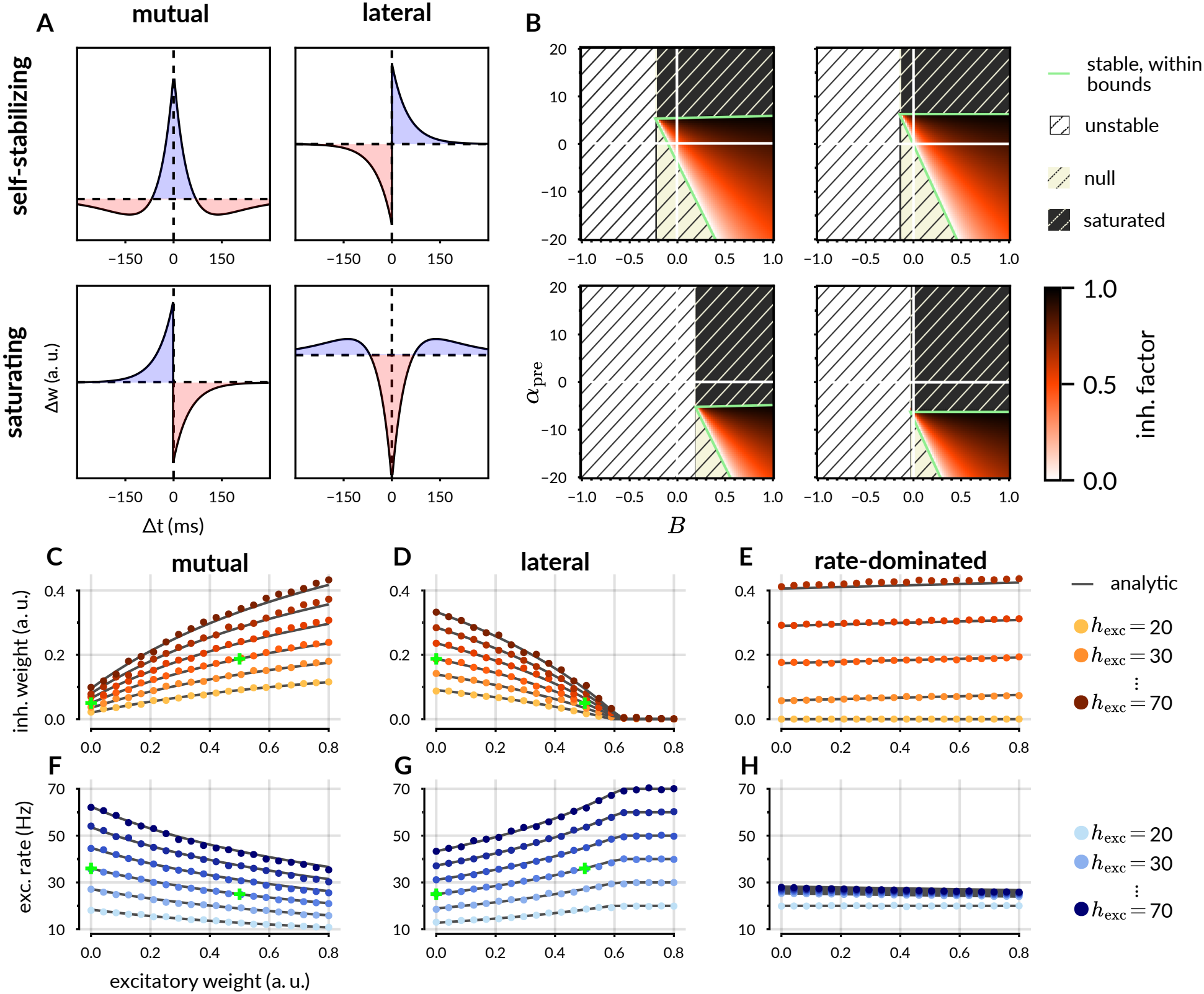
Parametric study of different shapes of iSTDP kernel in the reduced 2D model. **A.**Representation of four possible shapes of pairwise iSTDP interaction kernel, labeled by whether the synaptic weight converges to a finite interior fixed point (“self-stabilizing”) or diverges to the weight boundaries (“saturating”) in the covariance-dominated regime (i.e. with near-zero rate-dependent terms). **B**. Parametric study based on the analytic solution for each of the four iSTDP kernels, varying *α*_pre_ and the net area under the iSTDP kernel, *B* (see Eq. 1). The color map represents the level of inhibition, where 1 indicates that the excitatory neuron is completely silenced, and 0 indicates no inhibition, so that the excitatory firing rate matches the input current. **C**,**D**,**E**. Final inhibitory weight as a function of the incoming excitatory weight for three different iSTDP rules. Dots are numerical simulations, gray lines are analytic results (Methods, Eq. 11), color shades indicate different input levels. **C**. Results for the symmetric, self-stabilizing rule. The green dots represent the same conditions shown in Fig. 1F. **D**. Results for the antisymmetric, self-stabilizing rule. The green dots represent the same conditions shown in Fig. 1G. **E**. Results for the symmetric rule in a rate-dominated regime. **F**,**G**,**H**. Corresponding plots showing the rate of the excitatory neuron after convergence.

Our analysis found that both symmetric and antisymmetric kernels yield stable fixed-points in the covariance-dominated regime (Fig. 2A,B, top row, and Supporting Information Fig. S2C,D). These kernels, which we term “self-stabilizing”, exhibit robustness to parameter fluctuations and reliably converge to a finite inhibitory weight. In contrast, the inverted antisymmetric and the inverted symmetric kernels yield only unstable fixed points in the covariance-dominated regime ; consequently, the synaptic weight does not converge to an intermediate structured solution, but instead diverges toward the boundary values. We categorize these as “saturating” kernels to reflect this runaway weight saturation (Fig. 2A,B, bottom panels, and Fig. S2E,F in Supporting Information). Saturating kernels can still reach stability with strong rate-dependent terms, which reduce the overall influence of the kernel shape on the final weight.

This difference in stability arises from how the kernels interact with changes in pre- and postsynaptic correlations. Consider a synapse operating under a covariance-dominated plasticity rule at its fixed point. At equilibrium, potentiation and depression events balance exactly. A perturbation that strengthens the inhibitory weight leads to a reduction in pre-post firing events due to the increased inhibition. For self-stabilizing kernels, which have a positive right branch (corresponding to pre-post events), this reduction in correlation reduces the occurrence of synaptic potentiation events, shifting the balance in favor of synaptic depression. The synapse therefore depresses, eventually compensating the initial perturbation from equilibrium. Conversely, saturating kernels show the opposite effect: since their kernel is negative in the pre-post region, a reduction of pre-post correlation results in fewer synaptic depression events, so the initial balance is further shifted in favor of synaptic potentiation, amplifying the perturbation and driving the synaptic weight toward its imposed upper bound.

To further characterize the self-stabilizing rules, we extended our analysis beyond the U and M circuit configurations presented in Fig. 1, exploring a broader range of excitatory-to-inhibitory weights *w*_exc_ and input currents to the excitatory neuron. For the symmetric kernel, the learned inhibitory weight increases with *w*_exc_ (Fig. 2C), while for the antisymmetric kernel it decreases (Fig. 2D). Increasing the excitatory external input leads to stronger inhibitory weights in both cases (Fig. 2C,D), however the rate of the excitatory neuron still scales linearly with the input strength (Fig. 2F,G). In contrast, the rate-homeostatic rule exhibits a minimal dependence on *w*_exc_ (Fig. 2E) and a very tight control on the excitatory rate (Fig. 2H), which restricts the dynamic range of the excitatory neuron, limiting its ability to respond to changes in input. The E/I interactions in the two-neuron circuit allow for the weights to converge without saturation, differently from a two-neuron circuit with only excitatory neurons (Babadi and Abbott 2013).

In summary, our analysis of the reduced E/I circuit reveals a spectrum of iSTDP rules with distinct outcomes on connectivity and dynamics. On the one hand, covariance-dominated rules with self-stabilizing kernels enable structured stabilization (stable activity and specific connectivity motifs). As a consequence, they selectively reinforce connectivity motifs based on pre- and post-synaptic correlations, and allow a broader range of excitatory firing rates. On the other hand, rate-dominated rules prioritize homeostatic control of excitatory activity over the formation of specific connectivity motifs. Thus, these rules ensure robust dynamical stability, potentially at the cost of computational flexibility. These results demonstrate a trade-off between the emergence of specific excitatory-inhibitory motifs and the tight regulation of excitatory rates in the two-neuron circuit.

### 2.3 Structured stabilization in randomly connected RNNs

How do our results on structured stabilization from two-neuron E/I circuits translate to large recurrent networks? We simulated a randomly connected RNN of conductance-based leaky integrate-and-fire (LIF) neurons, with 900 excitatory and 100 inhibitory units. We modeled the excitatory connections (exc-to-exc and exc-to-inh) as fixed and sparse, while the inh-to-exc connections were modeled as all-to-all and initially small, but subject to iSTDP (Table 2). In this case the firing rate of each inhibitory neuron was not fixed as in the two-neuron E/I circuit, but was determined by the total amount of input into each neuron from its presynaptic partners, without any external constraint.

**Table 2:**
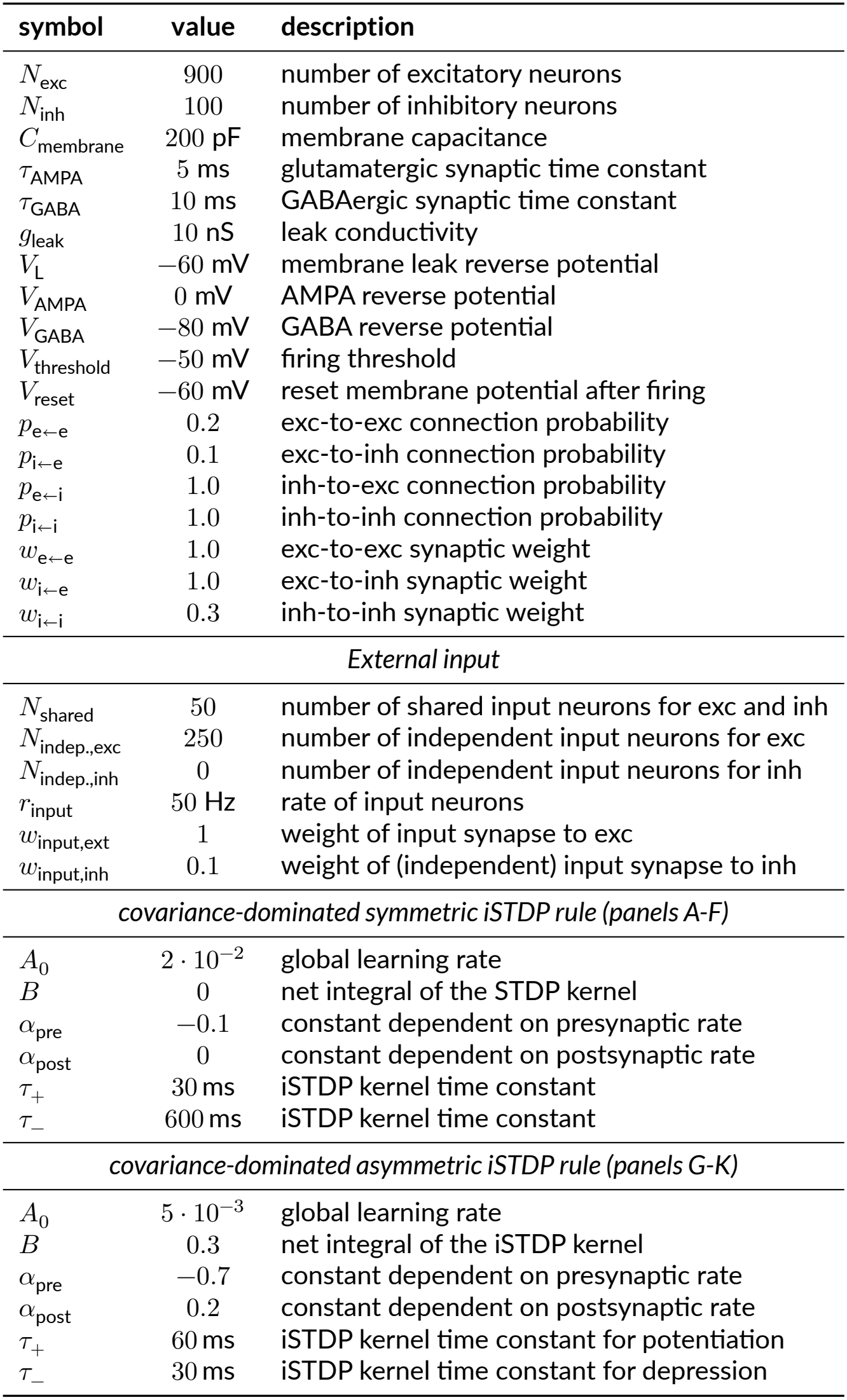
Numerical parameters for the random RNN in Fig.3.

| symbol | value | description |
|---|---|---|
| N<sub>exc</sub> | 900 | number of excitatory neurons |
| N<sub>inh</sub> | 100 | number of inhibitory neurons |
| C<sub>membrane</sub> | 200 pF | membrane capacitance |
| τ<sub>AMPA</sub> | 5 ms | glutamatergic synaptic time constant |
| τ<sub>GABA</sub> | 10 ms | GABAergic synaptic time constant |
| g<sub>leak</sub> | 10 nS | leak conductivity |
| V<sub>L</sub> | −60 mV | membrane leak reverse potential |
| V<sub>AMPA</sub> | 0 mV | AMPA reverse potential |
| V<sub>GABA</sub> | −80 mV | GABA reverse potential |
| V<sub>threshold</sub> | −50 mV | firing threshold |
| V<sub>reset</sub> | −60 mV | reset membrane potential after firing |
| p<sub>e←e</sub> | 0.2 | exc-to-exc connection probability |
| p<sub>i←e</sub> | 0.1 | exc-to-inh connection probability |
| p<sub>e←i</sub> | 1.0 | inh-to-exc connection probability |
| p<sub>i←i</sub> | 1.0 | inh-to-inh connection probability |
| w<sub>e←e</sub> | 1.0 | exc-to-exc synaptic weight |
| w<sub>i←e</sub> | 1.0 | exc-to-inh synaptic weight |
| w<sub>i←i</sub> | 0.3 | inh-to-inh synaptic weight |
| *External input* |  |  |
| N<sub>shared</sub> | 50 | number of shared input neurons for exc and inh |
| N<sub>indep.,exc</sub> | 250 | number of independent input neurons for exc |
| N<sub>indep.,inh</sub> | 0 | number of independent input neurons for inh |
| r<sub>input</sub> | 50 Hz | rate of input neurons |
| w<sub>input,ext</sub> | 1 | weight of input synapse to exc |
| w<sub>input,inh</sub> | 0.1 | weight of (independent) input synapse to inh |
| *covariance-dominated symmetric iSTDP rule (panels A-F)* |  |  |
| A<sub>0</sub> | 2 · 10<sup>−2</sup> | global learning rate |
| B | 0 | net integral of the STDP kernel |
| α<sub>pre</sub> | −0.1 | constant dependent on presynaptic rate |
| α<sub>post</sub> | 0 | constant dependent on postsynaptic rate |
| τ<sub>+</sub> | 30 ms | iSTDP kernel time constant |
| τ<sub>−</sub> | 600 ms | iSTDP kernel time constant |
| *covariance-dominated asymmetric iSTDP rule (panels G-K)* |  |  |
| A<sub>0</sub> | 5 · 10<sup>−3</sup> | global learning rate |
| B | 0.3 | net integral of the iSTDP kernel |
| α<sub>pre</sub> | −0.7 | constant dependent on presynaptic rate |
| α<sub>post</sub> | 0.2 | constant dependent on postsynaptic rate |
| τ<sub>+</sub> | 60 ms | iSTDP kernel time constant for potentiation |
| τ<sub>−</sub> | 30 ms | iSTDP kernel time constant for depression |

We first implemented a covariance-dominated plasticity rule with a symmetric kernel (Fig. 3A). Initially, the network produces a pathological high-firing regime, due to the low inhibitory weights. However, inhibitory plasticity quickly stabilizes the dynamics, generating an asynchronous irregular firing regime characteristic of balanced networks (Fig. 3B). To analyze the resulting inhibitory connectivity, we categorized inh-to-exc synaptic weights as either “mutual” or “unidirectional”. We called an inh-to-exc weight mutual when a reciprocal exc-to-inh connection exists between the same two neurons, reflecting a bidirectional link. Conversely, a weight is unidirectional in the absence of such excitatory reciprocal connection. Following the onset of plasticity, we observed that mutual inhibitory weights increase much more than unidirectional ones, with a portion of them reaching a saturation threshold, necessary for overall stability (Fig. 3C). This selectivity can be directly visualized through the matrix of synaptic weights after learning: the pattern of learned inh-to-exc weights mirrors the fixed exc-to-inh weights (Fig. 3E,F; see Supporting Information for full weight matrix).

**Figure 3:**
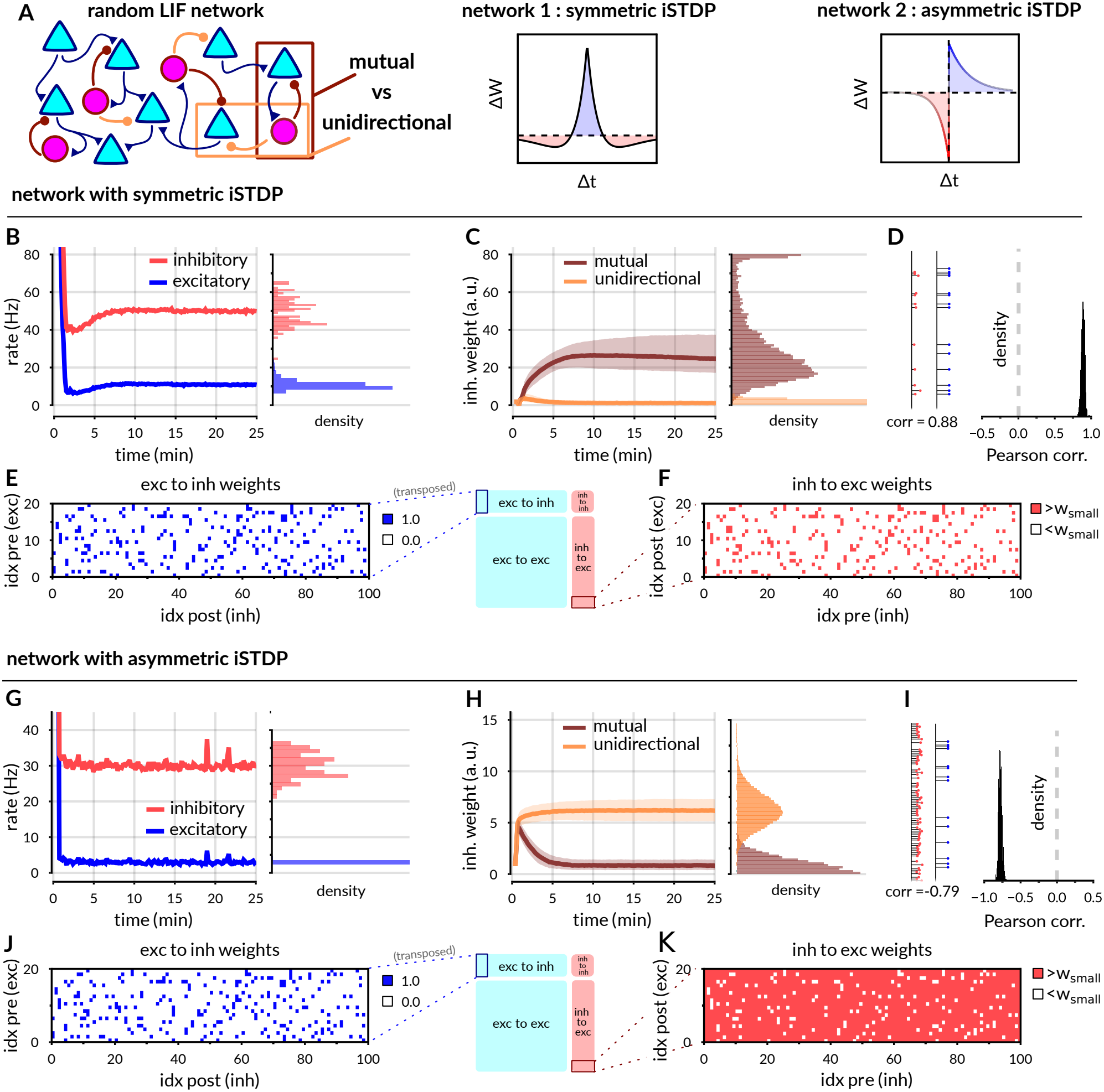
RNNs of conductance-based LIF neurons self-organize into specific E/I motifs under different iSTDP rules. **A** *left*. Representation of the random network with (*N*_exc_ = 900, *N*_inh_ = 100) spiking neurons. The inh-to-exc weights form all-to-all connections and are plastic, all other weights are fixed. Exc-to-exc and exc-to-inh connections are sparse. **A** *middle*. Pairwise interaction kernel of the symmetric, covariance-dominated iSTDP rule used in network 1 (panels B-F). **A** *right*. Kernel of the asymmetric, covariance-dominated iSTDP rule used in network 2 (panels G-K). **B.** Evolution of population mean firing rates over time, with final distribution of rates over the full populations. **C**. Evolution of inh-to-exc synaptic weights over time and final distribution. The weights are split into a “mutual” (brown) and a “unidirectional” (light orange) group. Only few weights reach the saturating value, set to 80. In panels **C** and **H**, thick lines show the median and shaded areas span the 25th to 75th percentiles of the synaptic weight distributions. **D**. Example of incoming (blue) and outgoing (red) weights from a single inhibitory neuron, with correlation between them, and distribution of correlations for the full inhibitory populations across 10 random network realizations. **E**. Random portion of the fixed and sparse exc-to-inh weights (the full matrix, of size 900×100 is shown in SI, Fig. S5). **F**. Portion of the all-to-all and plastic inh-to-exc connections after learning, with indices matching panel **E**. For clarity the matrix has been binarized, with threshold-value *w*_small_ defined as 0.1 times the 0.9-quantile of the full weight distribution. See SI, Fig. S5, for a continuous representation of synaptic weights. **G-K**. Equivalent plots in a second network with identical network dynamics and initial conditions, but using the asymmetric iSTDP (see also SI, Fig. S7). Parameters were determined through heuristic choices and coarse grid exploration (Methods, Section 4.5; Table 2).

Next, we investigated the impact of an asymmetric iSTDP rule within the same random RNN architecture and neural dynamics. As with the symmetric rule, this asymmetric iSTDP stabilizes the network dynamics (Fig. 3G). However, it promotes a markedly different connectivity pattern: purely lateral inhibition. In this configuration, inhibitory neurons strengthen their connections onto excitatory neurons lacking reciprocal connections back to them, thus forming exclusively unidirectional connections (Fig. 3H). This yields an inhibitory-to-excitatory connectivity pattern almost perfectly opposite to the previous configuration, so that existing exc-to-inh synapses carve gaps in the inh-to-exc connections (Fig. 3J,K). The asymmetric kernel tends to oppose synaptic growth in both pre-post and post-pre directions. A positive covariance at negative lags overlaps the depressive left side, while a negative covariance at positive lags overlaps the potentiating right side. Both pre-post and post-pre weights therefore contribute negatively to Δ*W*. Achieving robust lateral inhibition with the asymmetric iSTDP rule therefore requires the introduction of a small rate-dependent component (*B* = 0.3). This deviation from a purely covariance-dominated regime introduces characteristics of rate-homeostatic plasticity, such as rapid convergence to a narrow distribution of excitatory firing rates (Fig. 3G). Both the symmetric and the asymmetric rules converged to stable weight distributions, except for a very slow tendency for some mutual weights to reach either the upper bound (symmetric) or zero (asymmetric). These results are also robust to large variations in external input, and are compatible with network models where asynchronous irregular firing is generated intrinsically, without any external source of random noise (SI, Section S3 and Fig. S6; see also Mastrogiuseppe and Ostojic 2017).

In sum, our RNN simulations generalize the analytical predictions from the reduced E/I circuit model to larger networks of recurrently connected integrate-and-fire units. Symmetric iSTDP rules reinforce mutual inhibition, mirroring pre-existing excitatory connectivity, whereas asymmetric kernels favor lateral inhibition.

### 2.4 Response properties of a one-dimensional ring network

Having shown that different forms of iSTDP shape E/I connectivity in large-scale recurrent networks based on local correlations arising from direct connections, we next explored the functional consequences of structured stabilization in a recurrent network with a one-dimensional ring architecture. Ring networks are well-suited to model continuous attractor dynamics, supporting bumps of persistent activity (Amari 1977; Ben-Yishai et al. 1995; Khona and Fiete 2022). This makes them relevant for studying neural computations such as head-direction encoding (Xie et al. 2002; Kim et al. 2017; Kim et al. 2019; Petrucco et al. 2023) and orientation selectivity in the primary visual cortex (Goldberg et al. 2004).

We built a ring network of 800 excitatory and 200 inhibitory neurons arranged on a circle, with exc-to-exc and exc-to-inh weights decaying exponentially with the angular distance between neurons, following a rescaled Von Mises (Fig. 4A,B). The exc-to-PV projections have a broader spatial profile than the exc-to-exc and exc-to-SST projections; we also tested a network with identical spatial ranges for all outgoing excitatory connections (SI, Fig. S12). We further divided the inhibitory population into two subclasses, governed by distinct iSTDP rules, but sharing identical neural dynamics. Inspired by experimental results (Lagzi et al. 2021), one population of “PV” interneurons followed a symmetric kernel, and another population of “SST” interneurons followed an antisymmetric one, both rules operating in the covariance-dominated regime. Initially, we set both PV-to-exc and SST-to-exc synapses as all-to-all, weak, and unbiased (see Fig. 4A for schematics, and Methods Table 3 for numerical parameters).

**Table 3:** Numerical parameters for the one-dimensional ring model. The parameters that regulate neural dynamics are the same as in Table 2.

| symbol | value | description |
|---|---|---|
| *N*<sub>exc</sub> | 800 | number of excitatory neurons |
| *N*<sub>PV</sub> | 100 | number of PV interneurons |
| *N*<sub>SST</sub> | 100 | number of SST interneurons |
| sum *w*<sub>•←e</sub> | 145 | total incoming exc weights to exc, PV, SST |
| *k*<sub>e←e</sub> | 10 | width of exc-to-exc connections |
| *k*<sub>PV←e</sub> | 5 | width of exc-to-pv connections |
| *k*<sub>SST←e</sub> | 10 | width of exc-to-sst connections |
| *p*<sub>PV,SST←PV,SST</sub> | 0.9 | connection probability for PV-PV, SST-SST and PV-SST |
| *w*<sub>PV←PV</sub> | 2 | PV-to-PV synaptic weight |
| *w*<sub>SST←PV</sub> | 0.5 | PV-to-SST synaptic weight |
| *w*<sub>SST←SST</sub> | 2 | SST-to-SST synaptic weight |
| *w*<sub>PV←SST</sub> | 0.5 | SST-to-PV synaptic weight |
| *External input* |  |  |
| *h*<sub>in,exc</sub> | 1000 pA | constant input current to exc population |
| *N*<sub>input</sub> | 50 | number of independent input neurons |
| *w*<sub>input</sub> | 1.0 | weight of input synapse |
| *r*<sub>input,exc</sub> | 80 Hz | rate of input neurons to exc |
| *r*<sub>input,inh</sub> | 40 Hz | rate of input neurons to PV,SST |
| *symmetric iSTDP rule for the PV population* |  |  |
| *A*<sub>0</sub> | 2 · 10<sup>−3</sup> | global learning rate |
| *B* | 0 | net integral of the iSTDP kernel |
| *α*<sub>pre</sub> | −0.2 | constant dependent on presynaptic rate |
| *α*<sub>post</sub> | 1 | constant dependent on postsynaptic rate |
| *τ*<sub>+</sub> | 30 ms | STDP kernel time constant for potentiation |
| *τ*<sub>−</sub> | 600 ms | STDP kernel time constant for depression |
| *antisymmetric iSTDP rule for the SST population* |  |  |
| *A*<sub>0</sub> | 1 · 10<sup>−3</sup> | global learning rate |
| *B* | 0 | net integral of the iSTDP kernel |
| *α*<sub>pre</sub> | −0.2 | constant dependent on presynaptic rate |
| *α*<sub>post</sub> | 1 | constant dependent on postsynaptic rate |
| *τ*<sub>+</sub> | 30 ms | iSTDP kernel time constant for potentiation |
| *τ*<sub>−</sub> | 30 ms | iSTDP kernel time constant for depression |

**Figure 4:**
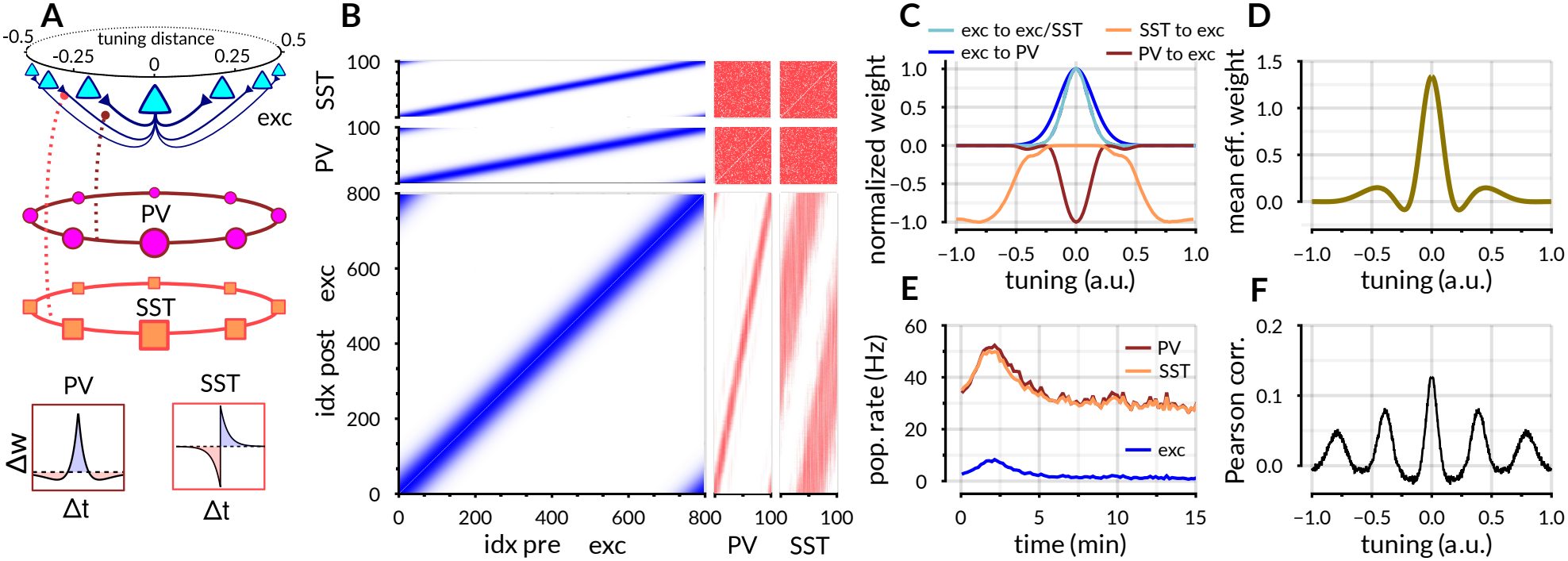
Inhibitory self-organization in a spiking RNN topologically organized as a one-dimensional ring. **A.**Representation of the model. 800 excitatory neurons are arranged in a one-dimensional ring topology, connected to each other and to their inhibitory counterparts with weights that depend on their distance in tuning space (Methods, Eq. 15). 100 “PV” inhibitory neurons connect all-to-all to the excitatory neurons with plasticity following a symmetric iSTDP rule, 100 “SST” neurons connect all-to-all to excitatory neurons with plasticity following an antisymmetric iSTDP. For both rules *B* = 0, *α*_pre_ = −0.1, *α*_post_ = 0. **B**. Full weight matrix after weight convergence. The excitatory weights (blue) are fixed, the inh-to-exc weights (red, lower right) are determined by plasticity. See SI, Fig. S8 for full weight matrix. **C**. Profile of the average outgoing weights as a function of tuning distance. For convenience, the profiles are normalized so that the maximum is either 1 or −1. **D**. Average effective interaction profile between the excitatory neurons, as a function of tuning (see Methods). **E**. Evolution of mean firing rate over time. **F**. Pearson correlation during spontaneous activity between an excitatory neuron and its neighbors, ordered by tuning distance and averaged over the entire population.

As the network evolves under the influence of the two plasticity rules, the inhibitory populations develop specialized roles. PV interneurons, governed by symmetric iSTDP, form strong mutual connections with excitatory neurons, mirroring the distance-dependent excitatory connectivity (Fig. 4B and Fig. 4C, red). Conversely, SST interneurons, driven by antisymmetric iSTDP, establish lateral inhibition across the ring (Fig. 4B and Fig. 4C, orange). This structured stabilization, with both localized and broader components, raises the possibility of nontrivial effective interactions between excitatory neurons, potentially mediated by the different roles of the PV and SST interneurons.

To estimate the effective interaction profile between excitatory neurons, we used linear response theory, equivalent to a Taylor expansion of the network dynamics around its operating point (Sadeh and Clopath 2020, see Methods for details). We found an effective connectivity profile resembling a Mexican hat, with short-range excitation directly inherited from exc-to-exc connectivity and a broader inhibitory surround resulting from the combined action of PV and SST interneurons (Fig. 4D). This effective connectivity profile motivated us to explore the resulting network dynamics and its response to external stimuli. We examined the emergent spontaneous activity and observed a complex spatial correlation structure, with periodic oscillations in correlations over distance (Fig. 4F, see Supporting Information Fig. S9 for full correlation matrix). This emergent dynamics is reminiscent of the spatial correlation maps observed in the developing cortex of ferrets, and is considered a signature of modular cortical organization (Smith et al. 2018; Powell et al. 2024). With identical spatial ranges for all outgoing excitatory connections, spatially structured correlations persist but decay over a shorter distance (SI, Fig. S12E; compare with Fig. 4F).

After identifying aspects of emergent cortical dynamics in our network consistent with early developmental patterns, we investigated whether it also displays other fundamental computational properties classically measured in the visual system. A well-established nonlinear phenomenon in the visual cortex is surround suppression, in which gradually increasing the stimulus size initially amplifies the neuron’s response up to a peak, but then leads to reduced activation (DeAngelis et al. 1994; Cavanaugh et al. 2002). After inhibitory plasticity self-organized network connectivity, we drove the network with patterned stimuli in the absence of subsequent plasticity. We delivered inputs of different strengths and sizes in the form of excitatory spike trains (Fig. 5A). Each stimulus was presented 100 times in randomized order, with a 0.5 s presentation followed by a 0.5 s blank. We found that larger stimuli elicited lower peak firing rates in their center region, while the response of neurons with receptive field closer to the stimulus’ edge was larger (Fig. 5A, raster plot in SI, Fig. S9). Consequently, the center neuron’s response as a function of stimulus size exhibited a classic surround suppression profile (Fig. 5B, DeAngelis et al. 1994).

**Figure 5:**
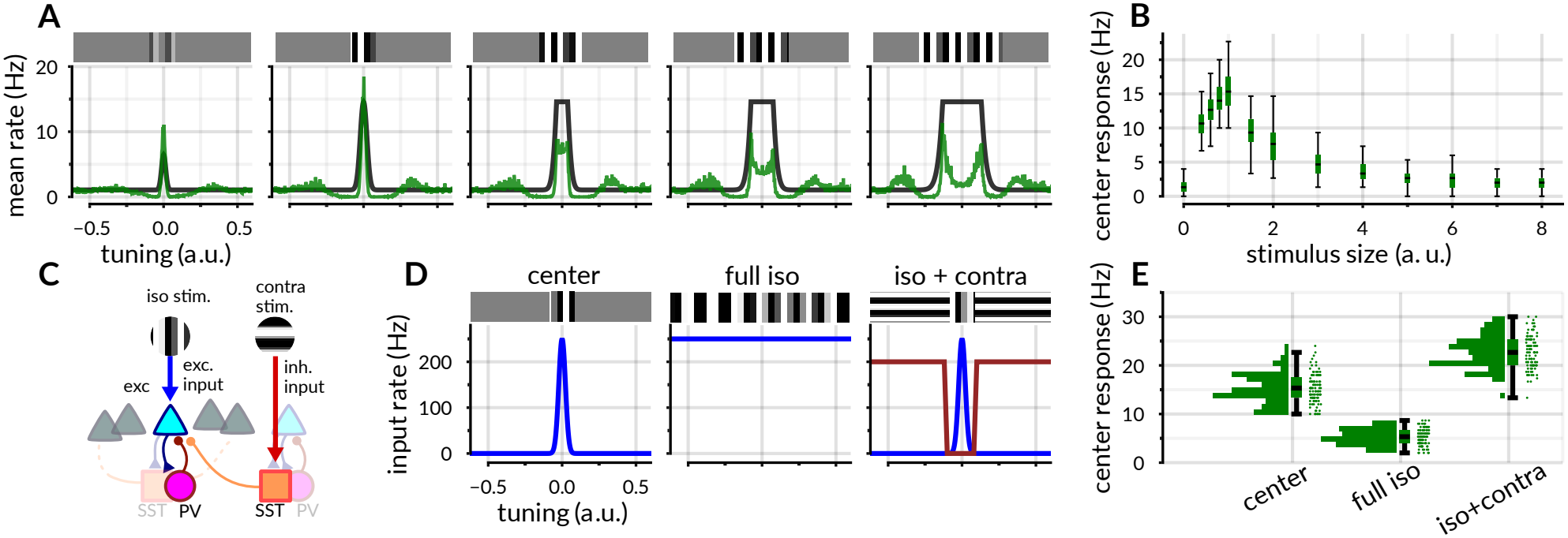
Response properties of the one-dimensional ring network. **A.**Representation of the stimuli (black) and of the network’s response (green), as a function of the neuron’s location relative to the stimulus center, averaged over 100 trials. **B**. Response to the center neuron only as a function of stimulus size. Boxes span the interquartile range (25th–75th percentiles), with the center line indicating the median and whiskers extending to the most extreme observations within 1.5 times the interquartile range. **C**. Schematics of the *iso-contra* stimulation protocol. The *iso* direction corresponds to excitatory inputs to excitatory neurons with matching receptive field. The *contra* stimulus instead provides inhibitory input to the SST population. **D**. Profiles of the center, full *iso* and *iso+contra* stimuli to the network. The blue line represents the excitatory input associated to the *iso* direction, the red line is the inhibitory input (i.e. affecting GABA conductance) associated to the *contra* direction. **E**. Distribution of center neuron responses in 100 independent trials for the three stimuli. Box-and-whisker conventions are as in panel B.

We carried out three control experiments to disentangle the contributions of the two interneuron populations with different plasticity rules to the dynamics of the ring network. A single inhibitory population with rate-dominated iSTDP can stabilize the system, although the emergent inh-to-exc connectivity remains largely unstructured over the simulated learning period (Fig. S10B). This network can still produce surround suppression, however the edge effect is reversed, with neurons on the stimulus edge firing at a lower rate compared to neurons in the center (Fig. S10C, note the convex shape of the curve for large stimuli). Moreover, the correlations do not exhibit any spatial structure. When considering networks with a single covariance-dominated PV-like (symmetric iSTDP kernel) or SST-like (asymmetric iSTDP kernel) plasticity rule, we found only pathological solutions that required careful parameter tuning. The PV-only network displayed pathological population-wide synchrony (Fig. S11E), while the SST-only network produced strongly clustered runaway excitation (Fig. S11J). These control simulations demonstrate that the two interneuron populations with different plasticity rules act synergistically to establish excitatory firing rate stability and specific E/I motifs.

We also investigated whether our model could reproduce contextual facilitating effects, for example when a stimulus in the preferred orientation is surrounded by an orthogonal stimulus (Sillito and Jones 1996). These effects are mediated by a third subclass of interneurons, the vasoactive intestinal peptide (VIP)-expressing cells, that inhibit the SST population and thereby disinhibit excitatory neurons (Keller et al. 2020). We thus modeled the *iso*-oriented stimulus as excitatory spike trains delivered to the excitatory neurons, and introduced a *contra*-oriented stimulus as inhibitory inputs from VIP interneurons to the SST population at the corresponding location (Fig. 5C). We then compared the center-neuron response in three conditions: (1) an *iso*-oriented stimulus restricted to the neuron’s receptive field; (2) a broader *iso*-oriented stimulus; (3) an *iso* stimulus in the receptive field combined with a *contra* stimulus in the distal regions (Fig. 5D). In agreement with experimental results (Keller et al. 2020), the distal *contra* stimulus disinhibits the center neuron and facilitates its response in the *iso+contra* stimulus conditions (Fig. 5E).

These results demonstrate how structured stabilization shaped by the interplay of symmetric and antisymmetric iSTDP rules can give rise to network connectivity which generates activity consistent with both suppressive and facilitating contextual effects in the primary visual cortex, and to long-range correlation patterns consistent with spontaneous activity in the developing sensory cortex.

## 3 Discussion

In this work, we have shown that a family of inhibitory STDP rules can simultaneously stabilize excitation at the circuit level, and drive the formation of specific connectivity patterns among excitatory and inhibitory neurons in the form of E/I motifs. In the absence of correlated external inputs, or training signals, circuits self-organize purely through their spontaneous internal dynamics. We found that the shape of the iSTDP rule selectively reinforces different E/I motifs—blanket inhibition, reciprocal E/I connectivity and lateral E/I connectivity—based on the correlations present in the network.

Prior computational studies addressing the self-organized formation of motifs in recurrent networks through STDP mostly focused on exc-to-exc plasticity (Ocker et al. 2015; Ravid Tannenbaum and Burak 2016; Triplett et al. 2018; Montangie et al. 2020; Manz and Memmesheimer 2023; Cusseddu et al. 2026). Previous work on iSTDP has been largely restricted to feedforward circuits and rules dominated by either firing rates or covariance (Kleberg et al. 2014; Luz and Shamir 2014; Akil et al. 2021; Vignoud and Robert 2022). In contrast, our approach combines both: recurrence and iSTDP. This combination makes it possible to characterize how different timing-based rules at inhibitory synapses shape excitatory-inhibitory connectivity in recurrent circuits, and how the resulting E/I motifs influence circuit dynamics, including rate stabilization, surround modulation, and the structure of spontaneous activity.

We implemented a generalized iSTDP framework where individual contributions from spike pairs additively contribute to synaptic change, as in many models of excitatory synaptic plasticity. While many other factors may also contribute to inhibitory plasticity, including multiplicative effects (Morrison et al. 2007; Kunkel et al. 2011), synaptic location (Agnes and Vogels 2024) or plasticity on behavioral timescales (Gonzalez et al. 2023), we found that additive iSTDP can still generate many interesting insights about E/I connectivity in recurrent networks. Our iSTDP framework combines both rate-dependent and covariance-dependent components in inducing inhibitory plasticity. The rate-dependent terms can stabilize neural activity through homeostatic regulation of firing rates, while the covariance-dependent terms can both stabilize activity and support the emergence of specific E/I motifs, motivated by recent experimental evidence (Znamenskiy et al. 2024). Consequently, our framework provides deeper insight into both the regulatory and structural properties of generalized iSTDP rules, expanding our understanding of plasticity in closed-loop recurrent circuits.

One strength of our framework is that it clarifies how the iSTDP kernel interacts with other plasticity parameters. For example, in the rate-dominated regime synaptic dynamics are primarily governed by firing rates and the net area of the iSTDP kernel, although correlations can still shape connectivity on a slower timescale than rate stabilization (Lagzi and Fairhall 2024) (see Supporting Information, Section S2). In the covariance-dominated regime, the weight evolution depends on specific spike covariance between pre- and postsynaptic spiking and the iSTDP kernel shape. This sensitivity enables structure to emerge along with rate stabilization, while allowing for a wider distribution of firing rates. The covariance-dominated regime, however, can pose greater challenges for parameter tuning. Although our reduced-model analysis provides qualitative guidance, generalizing it to large conductance-based networks is not trivial. Parameters in these networks were therefore selected through heuristic choices and coarse grid exploration. A promising way forward is to employ numerical methods, for instance using simulation-based inference and deep learning, which have already been applied to optimize both excitatory and inhibitory STDP rules in recurrent circuits (Confavreux et al. 2024). Our analytical approach can guide and complement these efforts, helping clarify parameter degeneracies and simplifying the parameter search process.

Our network simulations focus on representative symmetric and antisymmetric kernels. Extending this analysis to intermediate kernels that combine symmetric and antisymmetric components (Yang and Doiron 2026) would help determine how motif selectivity and network stability vary between these cases. Testing such mixed rules across additional recurrent architectures would also clarify the conditions under which distinct inhibitory populations support the computations demonstrated here.

Here we derived analytic solutions for the two-neuron Poisson model, and relied on numerical simulations for larger networks. By contrast, a recent study showed that iSTDP can stabilize recurrent LIF networks and converge to stable solutions through a mean-field approach (Akil et al. 2021). To achieve analytical tractability, that work assumed a regime of weak interactions, where internally generated correlations become negligible as the size of the neural population grows, and plasticity was driven exclusively by externally-induced correlations. In our approach, finite network size, strong interactions, and a dominance of correlation-dependent terms together drive self-organization. It remains an open question whether these two analytic frameworks, based on different assumptions, can be integrated.

While our analytical approach primarily examined how small E/I circuits self-organize under different iSTDP rules, neural circuits can also form larger structures, including feedforward firing sequences and assemblies. Assemblies are groups of strongly connected neurons, often regarded as the fundamental building blocks of neural computations (Buzsáki 2010; Papadimitriou et al. 2020; Miehl et al. 2023). Previous work has examined how various iSTDP rules enable the learning, maintenance and self-repair of neural assemblies (Ocker and Doiron 2019; Lagzi et al. 2021; Bergoin et al. 2023; Bergoin et al. 2025). In particular, recent findings suggest that assemblies are formed and best preserved when inhibitory neurons are divided into subclasses that combine symmetric and antisymmetric iSTDP (Lagzi et al. 2021). Here, the first rule can maintain network stability while the second prevents assemblies from merging through lateral inhibition, as in our work. Related work has also found that a combination of positive symmetric (Hebbian) and negative symmetric (anti-Hebbian) covariance-based rules can form and maintain stable assemblies governed by similar principles (Bergoin et al. 2025). These results on assemblies agree with ours in underscoring the intuition that different rules serve different purposes: some promote E/I balance and internal network stability, while others introduce competition through lateral inhibition.

In this work we further combined multiple iSTDP rules in a network with a ring topology (Ben-Yishai et al. 1995) to demonstrate how their interaction stabilizes activity while also supporting specific network computations, such as contextual input modulation as observed in the visual cortex (DeAngelis et al. 1994; Cavanaugh et al. 2002), and long-range spatial correlations reminiscent of modular spontaneous activity observed in the developing cortex (Smith et al. 2018; Powell et al. 2024). The parametrization of iSTDP rules allowed us to explore the entire space of rules and link them to the emergence of specific E/I connectivity motifs beyond the choice of pre-set, fixed plasticity rules in existing work (Lagzi et al. 2021; Bergoin et al. 2023; Lagzi and Fairhall 2024; Bergoin et al. 2025).

In this ring-network implementation, the spatial profiles of the exc-to-exc and exc-to-inh connections and the assignment of distinct plasticity rules to the two inhibitory populations are fixed, whereas the inh-to-exc weights self-organize from initially weak, unstructured connections. Thus, while the local excitatory component of the Mexican-hat effective connectivity is prescribed, its inhibitory surround emerges through iSTDP. More broadly, these circuit simulations serve as an interpretable proof of principle for how distinct iSTDP rules can jointly shape connectivity and computation, rather than as a detailed model of realistic cortical PV/SST organization. As a further simplification, our circuit models lacked any externally patterned input during learning. It would be interesting to consider non-stationary inputs, such as input correlations varying in time and across neurons, or highly structured input such as assembly-activations or waves spreading across the circuit as observed during cortical development (Smith et al. 2018). Non-stationary input, jointly with excitatory STDP, might help paint a more complete picture of prolonged cortical development especially after the onset of sensory experience (Trägenap et al. 2025). Excitatory plasticity can enable circuits to learn new representations, refine current ones, and recover after lesions.

Overall, our findings highlight how local inhibitory learning rules can both stabilize global network dynamics and produce specific connectivity motifs in a self-organized manner in recurrent circuits, with implications for understanding neural development and learning under structured stimuli. While the specific cortical-like response patterns explored here rely on idealized architectural assumptions and their broader biological interpretation remains to be tested (e.g. across more generic recurrent architectures with multiple inhibitory populations, or against quantitative connectomics data), the underlying mechanistic principles linking iSTDP to emergent E/I motifs across different circuit structures provide a general foundation for future experimental and theoretical work.

## 4 Methods

### 4.1 General mean-field solution

We consider a class of pairwise iSTDP rules triggered by spiking events and purely based on spike-timing difference between pre- and postsynaptic neuron firing, defined by the following expression.

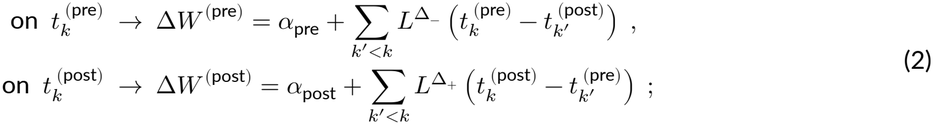

where 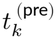 and 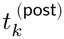 represent pre- and postsynaptic spike events, respectively. In our numerical simulations the instantaneous change of weight is further regulated by a global learning rate *A*_0_, as in Δ*W*^* (pre,post)^ = *A*_0_ Δ*W*^(pre,post)^. The scaling by *A*_0_ is omitted in our equations for simplicity. Each presynaptic spike triggers a weight change consisting of a fixed component, denoted *α*_pre_, and a pairwise interaction term. Similarly, each postsynaptic spike triggers a weight change with a fixed component *α*_post_ and a pairwise interaction term. The pairwise interaction terms capture the cumulative influence of joint spiking events, incorporating the contribution of all preceding spikes (*k*^′^ < *k*) weighted by a kernel 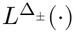 that depends on the time differences between the spikes. The two functions 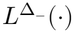 and 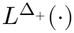 refer to the post-pre and the pre-post pairwise interaction, and are merged in a single function, known as the iSTDP pairwise interaction kernel, as follows:

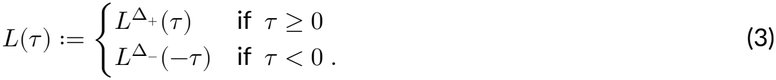

By convention, the *τ* > 0 branch of the curve represents pre-post interactions (presynaptic spike preceding the postsynaptic spike), while the *τ* < 0 branch represents post-pre interactions (postsynaptic spike preceding the presynaptic spike). In our framework, we consider *L*(*τ*) to be independent of *α*_pre_ and *α*_post_. This is in contrast to previous formulations that incorporate these terms into the kernel representation (e.g., Vogels et al. 2011 and Confavreux et al. 2024). We assume that synaptic plasticity operates on a much slower timescale than neural dynamics, allowing us to treat synaptic weights as constant when calculating first- and second-order firing statistics (Kempter et al. 1999). We further assume variability in the spike-train statistics, so that it can be effectively described as a stochastic process. Thus, we can write the evolution of the mean synaptic weights as:

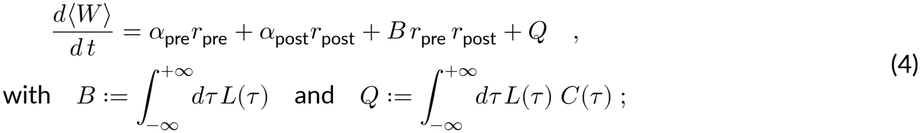

where *B* is the integral of the iSTDP kernel, and *Q* is the only term that depends on the pairwise covariance density between pre and post-synaptic spike trains, here denoted *C*(*τ*). For a complete derivation see Supporting Information or Kempter et al. (1999) and Luz and Shamir (2014).

### 4.2 Expressions for the symmetric and antisymmetric exponential kernel

Eq. 4 can be applied to any form of iSTDP kernel, including, e.g., kernels with delayed interactions (Babadi and Abbott 2013). Here, we focus on two specific exponentially decaying shapes: symmetric and antisymmetric, defined as follows:

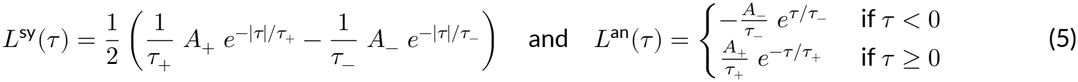

With this parametrization, *A*_+_ = *A*_−_ implies *B* = 0, corresponding to covariance-dominated rules, whereas for *A*_−_ = 0 we have *B* = *A*_+_, a rate-dominated regime. This choice of *L*(*τ*) allows us to keep track of the entire spiking history through activity traces updated at every spike time and decaying in time with the same constants as the iSTDP kernel. These traces will determine the amount of potentiation (+) or depression (−) triggered by each spike. For the antisymmetric rule, the presynaptic and postsynaptic traces are respectively defined as follows:

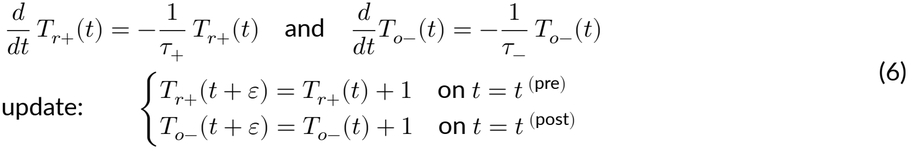

The symmetric rule requires, in addition to *T*_*r*+_(*t*) and *T*_*o*−_(*t*), the traces *T*_*r*−_(*t*) and *T*_*o*+_(*t*), defined analogously. The implementation of the asymmetric iSTDP rule is then given by:

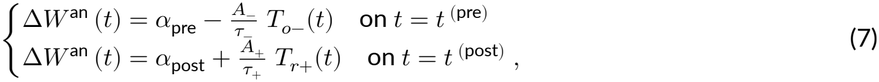

and for the symmetric rule:

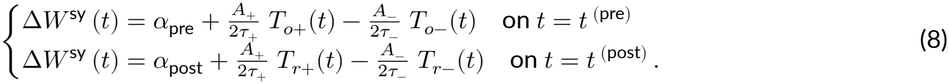

### 4.3 Low impact of correlations in rate-dominated rules

To better understand the role of the net area *B* of the iSTDP kernel in the weight dynamics (Eq. 4), and how it determines whether a rule is rate-dominated or covariance-dominated, we will consider here a simplified scenario, where *L*(*τ*) prescribes a +1 increase in weight for coincident pre-post spikes and no increase otherwise. This choice yields *B* = *A*_+_ = 1. Consider now a pre and a post-synaptic neuron that fire with Poisson statistics at rates *r*_pre_ = *r*_post_ = *r* = 10 Hz. The term *B r*_pre_ *r*_post_ in Eq. 4 takes a value of 100 s^−2^ regardless of the correlation between the two neurons. But how does it compare to the covariance term *Q* in the presence of strong correlations between pre and post-synaptic activity? We can consider a regime of (unnaturally) high correlations, such that half of the spikes of the two neurons are synchronous (Pearson correlation 0.5). The term *Q*, corresponding to the spike-count covariance, is 5 s^−2^. So even for very strong correlations *Q* is 20 times smaller than *B r*_pre_ *r*_post_. This illustrates that in regimes with strong *B*, the weight dynamics in Eq. 4 are primarily governed by first-order statistics, with — at best — second-order contributions from correlations in spiking activity.

### 4.4 The two-dimensional reduced model

Finding a general solution for Eq. 4 can be challenging. For our purposes, we consider a simplified case. First, we model an excitatory and an inhibitory neuron as mutually interacting Poisson processes (Hawkes 1971; Ocker et al. 2017). Each time the presynaptic neuron spikes, the firing probability of the postsynaptic neuron increases (or decreases) proportionally to the excitatory (or inhibitory) post-synaptic potential EPSP (IPSP) kernel:

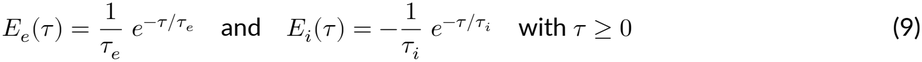

We further assume that the input to the inhibitory neuron *h*_inh_(*t*) is regulated homeostatically to keep its rate *r*_inh_constant. Namely,

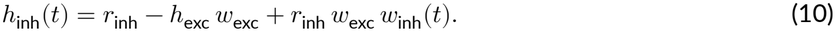

In those conditions, Eq. 4 has an approximate fixed-point closed-form solution for the inhibitory connection *w*_inh_:

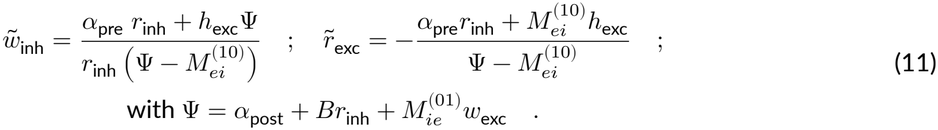

The coefficients 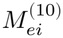 and 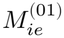 are found through the linear response theory (Hawkes 1971; Ocker et al. 2017), and depend on the iSTDP kernel and the EPSP / IPSP parameters (refer to the analytic derivation of the SI Text, Section S1.4).

For the symmetric iSTDP:

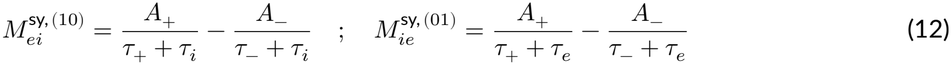

For the antisymmetric iSTDP instead:

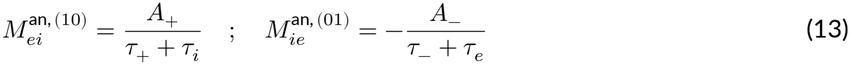

Numerical parameters are in Table 1.

### 4.5 Randomly connected recurrent neural network

We built a random network of conductance-based leaky integrate-and-fire (LIF) neurons, with 900 excitatory units and 100 inhibitory units. The membrane potential and the synaptic conductances obey the following dynamics:

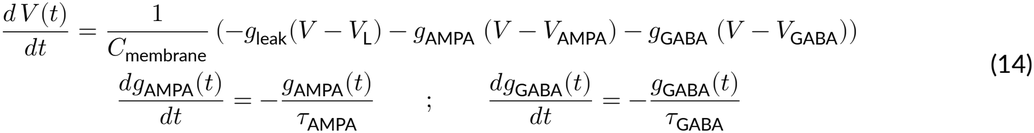

Each presynaptic excitatory (inhibitory) spike, both from the recurrent connections and from the input units, increased the AMPA (GABA) conductance, *g*_AMPA_ (*g*_GABA_). Each increase was a pulse-like update equal to the connection’s synaptic weight. We assigned a probability of connection for all non-plastic connections (exc-to-exc, exc-to-inh, and inh-to-inh). The inh-to-exc connectivity was all-to-all and plastic. Additional excitatory input units generated spikes as a Poisson process of fixed rate. A fraction of the input units was shared among all excitatory neurons, leading to a small correlated activity. The network displayed high robustness to variations in input strength (see SI, Fig. S13). Numerical parameters were determined through a mix of heuristics and coarse grid search (Table 2).

### 4.6 Ring model

The ring model was implemented with the same neural dynamics as the random network (Eq. 14). Here however both excitatory and inhibitory neurons received a baseline external input, independent for each unit. We used 800 excitatory units and 200 inhibitory units, further divided into a PV and an SST subclass. The synaptic weight between an excitatory neuron and its neighbors decayed with distance, following a bell-shaped curve modeled as a rescaled von Mises distribution:

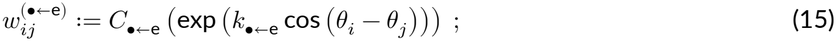

where *θ*_*i*_ and *θ*_*j*_ represent the locations of the post- and presynaptic neuron on the ring, with each population equally spaced in the [0, 2π) interval. The factor *k*_•+-e_ regulates the connection range. Finally the scaling factor *C*_•+-e_ is tuned so that the sum of incoming weights to either exc, PV or SST population is at a desired value sum *w*_•+-e_. The inh-to-inh connectivity is all-to-all except for autapses and fixed in time. We considered two distinct inhibitory populations with identical dynamics, but different iSTDP rules. The first population had a symmetric iSTDP kernel, the second an antisymmetric one. The input was different than in the previous model, with a constant external current in the excitatory population, and independent excitatory spike trains for all units.

To measure the network response to external stimuli (Fig. 5), we froze plasticity after learning, and delivered stimuli as uncorrelated spike trains. *Iso*-oriented stimuli took the form of excitatory spikes delivered to the exc neurons, *contra*-oriented stimuli were modeled as inhibitory spikes delivered to the SST neurons. Numerical parameters, determined through grid search, are shown in Table 3.

## 5 Software implementation and code availability

All simulations for the reduced models are based on custom code written in Julia (Bezanson et al. 2017, https://julialang.org), also available as separate toolbox. For the large-scale simulations we used Brian2 (Stimberg et al. 2019, https://briansimulator.org). Finally some of the analytic integrals were solved using Maple (https://maplesoft.com). The code is fully open-source and publicly available on GitHub under the Creative Commons Attribution 4.0 International (CC BY 4.0) License. https://github.com/comp-neural-circuits/structured-stabilization-in-recurrent-neural-circuits

## Acknowledgments

This project has received funding from the European Research Council (ERC) under the European Union’s Horizon Europe research and innovation programme (ERC CoG FeedbackCircuits, grant agreement no. 101170267), and under the Marie Skłodowska-Curie ITN SmartNets (grant agreement No. 860949) under the European Union’s Horizon 2020 research and innovation program, by the International Human Frontier Science Program Organization (RGP0062/2021), and by the Technical University of Munich (TUM). Thanks to the “Computation in Neural Circuits” group for their useful insights and discussions. Special thanks to Alessio Quaresima for his valuable feedback and for independently testing our results.

## Supporting Information for

### S1 Analytic results

#### S1.1 Mean-field equation for weight dynamics

In this section, we adapt the results introduced by Kempter et al. (1999) to our notation and our specific needs. We first describe a sufficiently general expression for the pairwise STDP rules. Given two neurons, one presynaptic and one postsynaptic, an excitatory or inhibitory STDP rule is a weight update triggered by either a pre- or a postsynaptic spike. The weight update solely depends on the time differences between the previous spike times, relative to the current spike time:

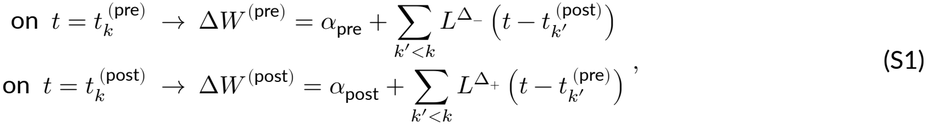

where *t*_*k*_^(pre)^ and *t*_*k*_^(post)^ represent pre- and postsynaptic spike events, respectively. We can merge the functions 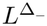 and 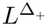 into a single STDP kernel defined as:

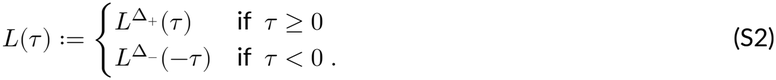

Therefore *τ* > 0 (the Δ_+_ branch) represents pre-post interactions (presynaptic spike preceding the postsynaptic spike), while *τ* < 0 (the Δ_−_ branch) represents post-pre interactions (postsynaptic spike preceding the presynaptic spike).

It is useful to express each spike train as a sum of Dirac delta functions centered at each spike time:

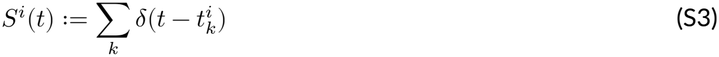

where *i* denotes the neuron index. With this notation, the number of spikes in the [*t*_*a*_, *t*_*b*_] interval is:

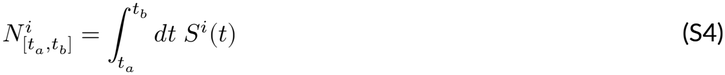

Assuming stationary conditions, we can therefore define the firing rate of neuron *i* as:

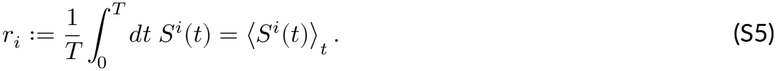

Using the definition in Eq. S3, we can then rewrite the pairwise interaction terms in Eq. S1 as:

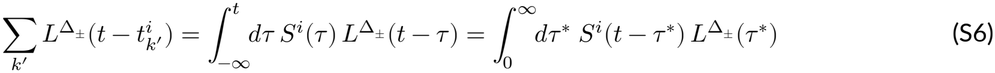

Where in the last step we performed the change of variable *τ* ← *t* − *τ* ^*^. This leads to the following expression for the instantaneous change in synaptic weights, numerically equivalent to Eq. S1:

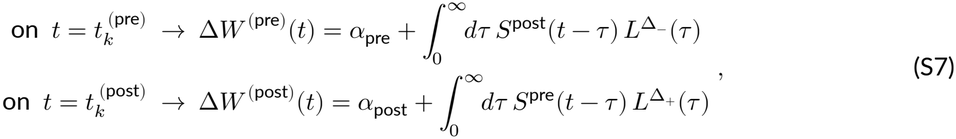

Now, using again the same definition of spike train in Eq. S3, an event-based update can be expressed as an integral over time over the pre- and post-synaptic spiking events. The total weight change occurring in an interval [*t*_*a*_, *t*_*b*_] is:

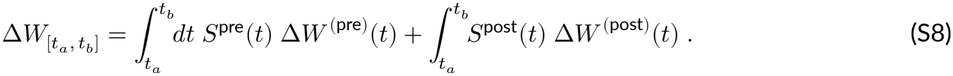

We can then express the average weight change per second, considering a time interval [0, *T*] large enough to ensure precise sampling in second-order firing statistics, but still small compared to the rate of changes in Δ*W*. Under this assumption of separation of timescales between neuronal dynamics and plasticity, we can substitute Eq. S7 into Eq. S8, and write:

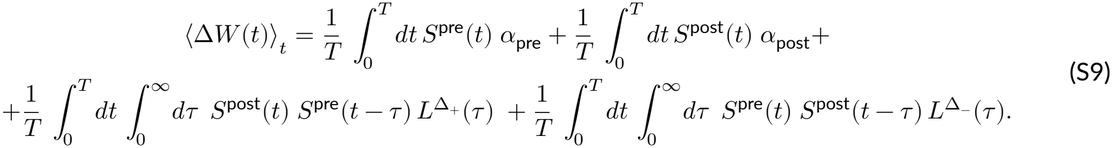

Since *α*_pre_ and *α*_post_ are constants, we can factor them out. The first two terms of the r.h.s. of Eq. S9 become:

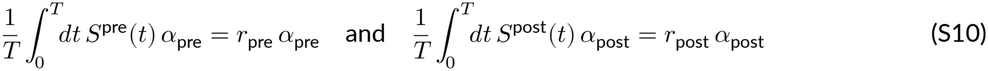

The pairwise interaction terms instead can be simplified as follows:

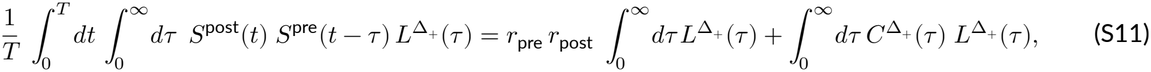

and similarly for the last term in Eq. S9. In the above equation,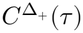 denotes the covariance density between the pre- and the postsynaptic neuron, following the convention that Δ_+_ represents the pre-post order and Δ_−_ the post-pre order:

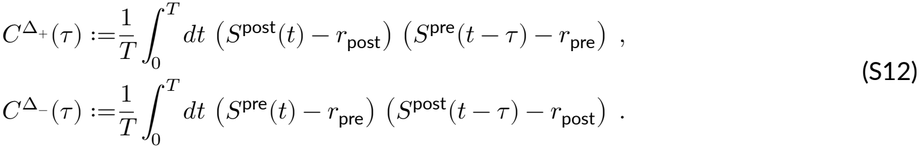

A change of variable gives 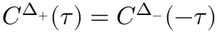. We therefore simplify the notation as follows: 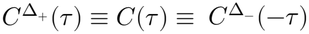. Note that this identity simply reverses the spike order, and does not imply that *C*(*τ*) = *C*(−*τ*). In our model, *C*(*τ*) is generally asymmetric (Fig. S1B-E). Substituting these definitions into Eq. S9 yields:

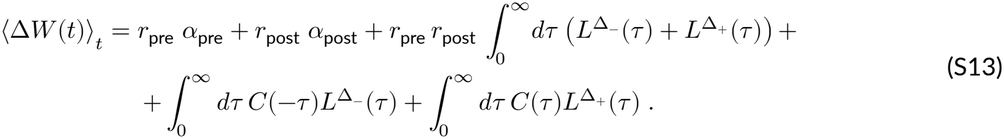

By the definition of the combined kernel in Eq. S2, the rate-dependent pairwise term becomes

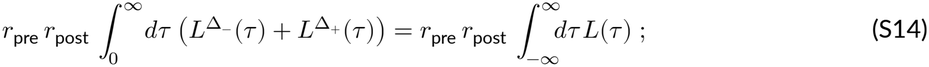

and we simplify through the following variable substitution:

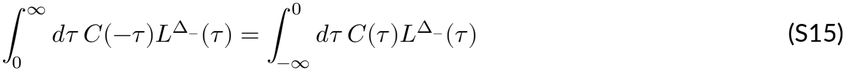

Finally, combining the post-pre and pre-post contributions, we have:

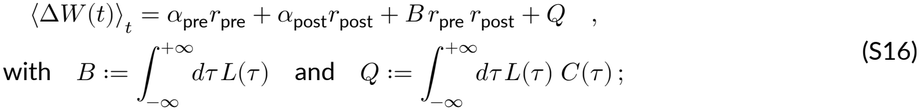

Corresponding to Eq. 1 in the main text.

### S1.2 Derivation of the closed-form solution for the E/I motif

In this section, we derive a closed-form solution for a reduced model of one inhibitory and one excitatory neuron (Eq. S16). We model the two neurons as mutually interacting Hawkes processes (Main Text, Fig. 1A, Hawkes 1971; Pernice et al. 2011). The excitatory unit receives a constant input current *h*_exc_, a plastic inhibitory connection *w*_inh_(*t*) and it projects a fixed excitatory connection *w*_exc_ to the inhibitory neuron. For simplicity, we assume that the inhibitory unit is kept at a constant rate *r*_inh_ by an adaptive external current *h*_inh(*t*)_ (10). In this framework, the average firing rates behave linearly, therefore, the excitatory rate can be written as:

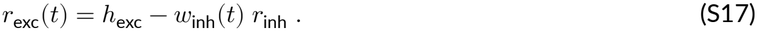

As we focus on the connection *w*_inh_(*t*) from the inhibitory to the excitatory neuron, we rename *r*_inh_ = *r*_pre_ and *r*_exc_ = *r*_post_ in Eq. S16 and we get:

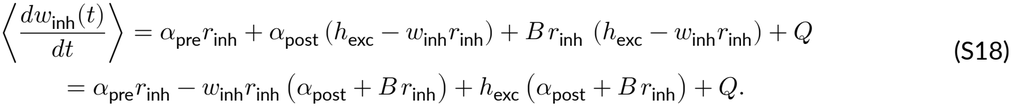

The only nontrivial quantity to compute is the term *Q*, as it requires the inhibitory-to-excitatory covariance density curve. To avoid ambiguities, we define the Fourier transform operator acting on a function *f* (*t*) and the convolution operator acting on two functions *f* (*t*) and *g*(*t*) as follows:

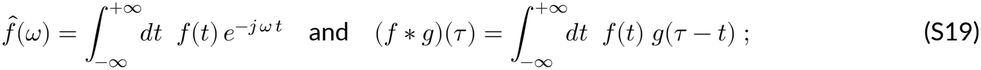

where *j* denotes the imaginary unit.

Note that the total integral *Q* of the product *C*(*τ*) *L*(*τ*) (as defined in Eq. S16) is equivalent to its Fourier transform evaluated at *ω* = 0. But for the inverse convolution theorem, the Fourier transform of the product *C*(*τ*) *L*(*τ*) corresponds to the convolution of the two transforms *Ĉ* (*ω*) and 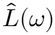. In equations:

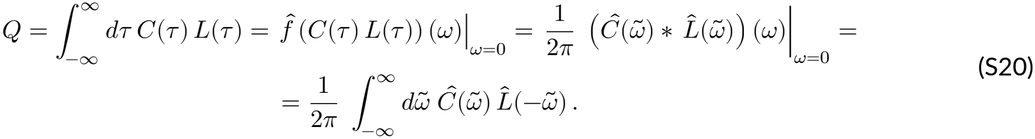

The advantage of re-defining *Q* in the Fourier space is that, for Hawkes processes, the covariance *Ĉ* (*ω*) can be expressed in a closed form as a function of network connectivity and neural activity (Hawkes 1971; Pernice et al. 2011; Trousdale et al. 2012; Jovanović et al. 2015). More specifically, given the matrix of post-synaptic potential kernels (Eq. 9 in Methods)

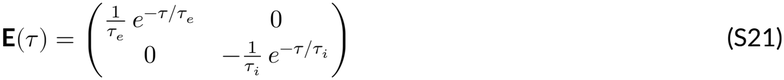

and the connectivity matrix

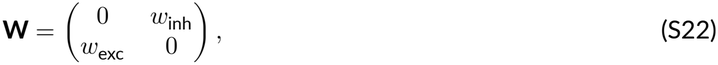

we can first write the vector of firing rates **r** = [*r*_exc_, *r*_inh_]^⊤^ as

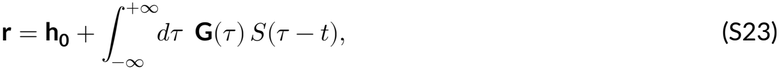

where **h**_**0**_ is the baseline input and **G**(*τ*) = **E**(*τ*) **W** is referred to as the matrix of “interaction kernels”. At the equilibrium state,

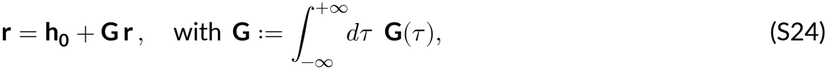

where the integral in the r.h.s. corresponds to the Fourier transform evaluated at zero, i.e., 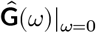. Equivalently, the rate vector can be written in an explicit form:

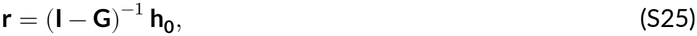

where **I** represents the identity matrix. Moreover, if we write the covariance matrix (cf. Eq. S12) as

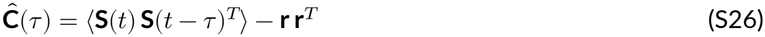

then we have the following expression in the Fourier domain, at the equilibrium state:

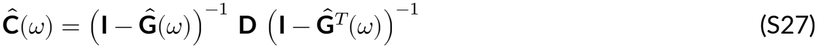

where **D** is a diagonal matrix with the firing rates on its diagonal. Finally, recall that, for a matrix **G** with spectral radius < 1,

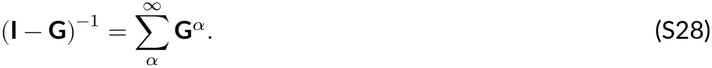

Therefore, the covariance matrix can be written as the following expansion

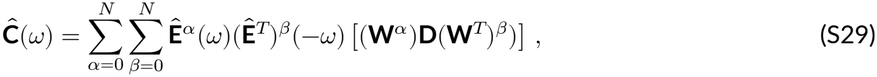

where the indices *α, β* indicate the number of connections between any spiking neuron in the network to the post- and to the presynaptic neuron, respectively. This captures the influence of each spike propagated in the network through existing synaptic connections on the correlated firing of the pre- and postsynaptic neurons. Within this formalism, in our reduced model we consider only pairwise interactions, that is, *α* + *β* = 1. The inhibitory-to-excitatory covariance is the (1, 2) element of 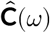. We can compute it as:

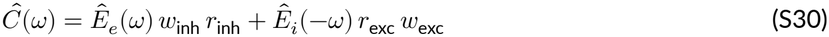

By combining Eq. S20 and Eq. S30, we get the following expression for *Q*:

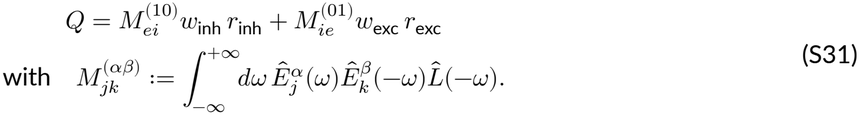

The 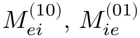 coefficients can be computed analytically depending on the iSTDP rule (see next section). Considering them as constants, we can then write Eq. S18 as:

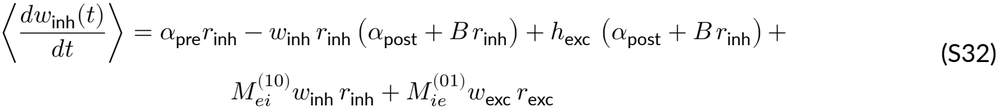

or equivalently,

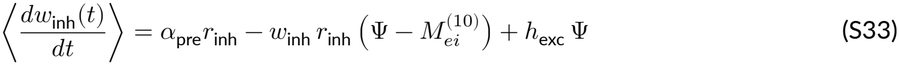

with 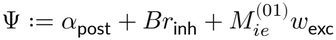.

We then find the stationary point (Eq. 11 in Methods)

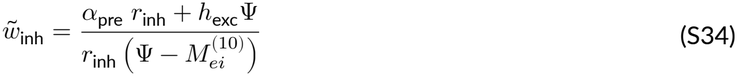

The equilibrium point 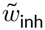 is a stable fixed point attractor, and positive, that is 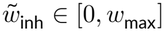, if the following conditions hold:

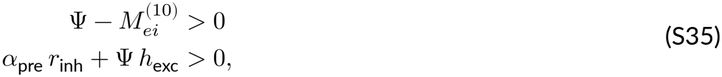

If a solution for 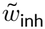 exists, we can also express the steady-state postsynaptic rate as follows:

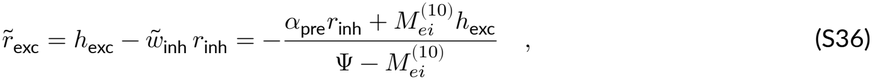

The solution is valid only for positive rates. This adds one more condition for a stable and positive solution. The constraints are therefore:

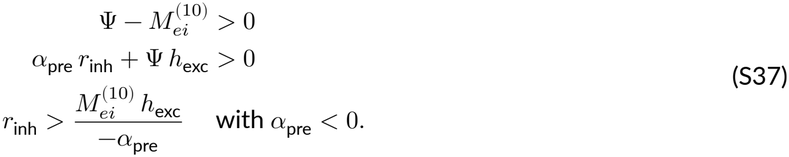

### S1.3 Circuit motif with unconstrained *r*_**inh**_

In the 2D E/I motif presented in the main text, the value of *r*_inh_ was fixed both for in the numerical simulations and in the analytic expression of the dynamics (Eq. 10). This led to a closed-form expression for the fixed point solution (Eq. 11 in Methods):

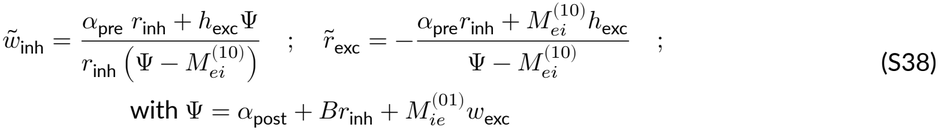

In this section, for completeness, we show how the condition of fixed *r*_inh_ is not required for a stable solution and can in fact be relaxed at the cost of simplicity in the analytic expression of the fixed-point solution.

Consider the model shown in Fig. S3A. Here both *r*_exc_ and *r*_inh_ are free parameters, given by the net input. Namely:

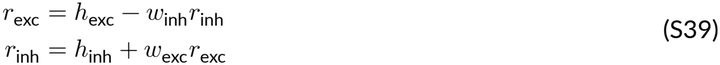

where we dropped the time dependence in the notation for simplicity. We can rewrite the equations above as:

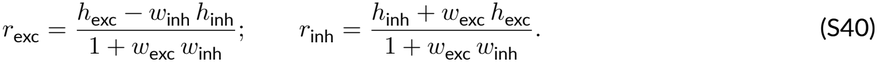

Then, plugging the terms above into (S32), we obtain:

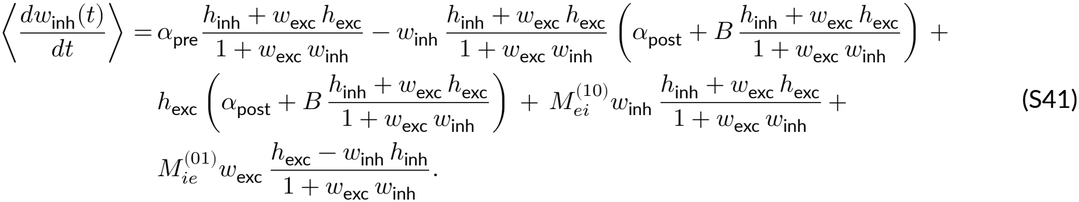

We can find a fixed point 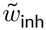 in the equation above by setting 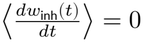 and multiplying by (1 + *w*_exc_ *w*_inh_)^2^. The equation becomes a quadratic equation for *w*_inh_:

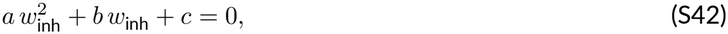

with

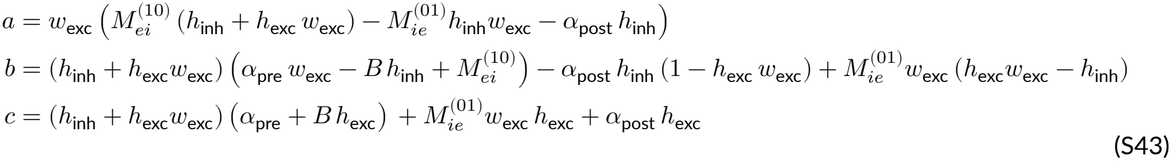

The solution is then given by

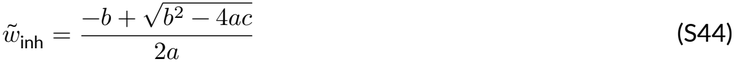

as long as *b*^2^ − 4*ac* ≥ 0.

Figure S3 shows analytic and numeric results for unidirectional and mutually-connected motif configurations for different iSTDP rules.

### S1.4 Derivation of first-order motif coefficients

To calculate the motif coefficients

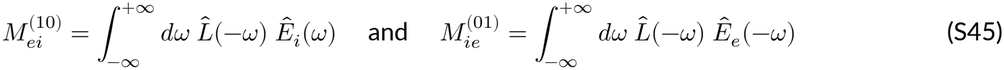

for the connection *W*, we need to compute the Fourier transform of the postsynaptic potential *E*_*e*_, *E*_*i*_ and the Fourier transform of the iSTDP kernel.

The Fourier transforms of the excitatory and inhibitory postsynaptic potential (Eq. 9 in Methods) are respectively given by:

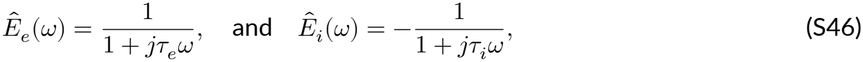

where *j* is the imaginary unit.

Consider first the Hebbian symmetric STDP kernel, defined via the function

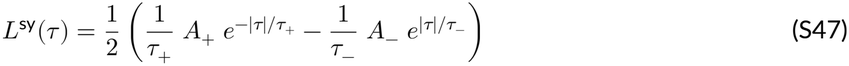

with *A*_+_, *A*_−_ > 0. The Fourier transform is given by

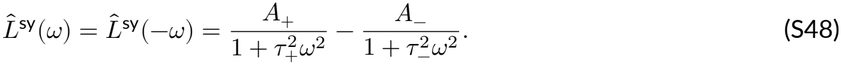

By substituting Eq. S46 and Eq. S48 in Eq. S45, we can calculate the first-order motif coefficients 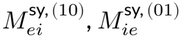:

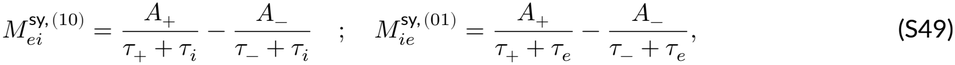

and therefore find an analytical solution for the integral *Q* in Eq. S31. Consider now the antisymmetric iSTDP rule, with kernel:

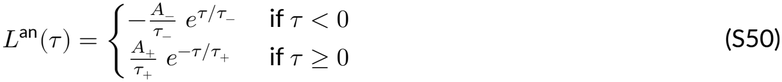

whose Fourier transform is given by

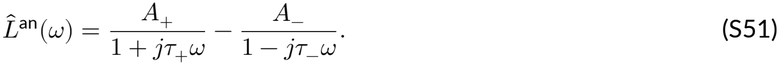

Again, we can calculate the first order motif coefficients 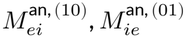 for the antisymmetric rule by substituting Eq. S46 and Eq. S51 in Eq. S45:

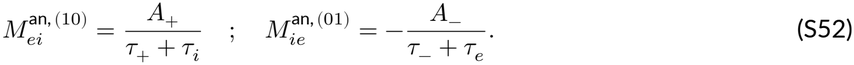

These expressions can be included into Eq. S34, leading to the final expression for the symmetric rule:

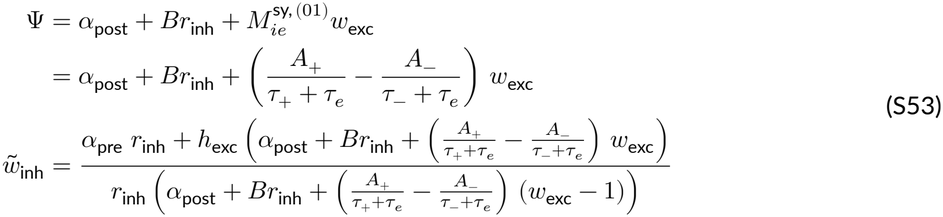

And to the final expression for the antisymmetric rule:

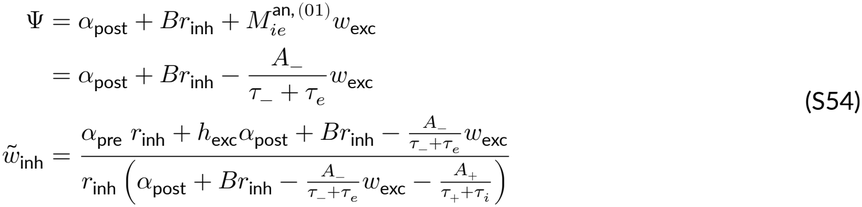

### S2 Correlation-dependent selectivity under rate-dominated versus covariance-dominated iSTDP rules

Previous work has shown that the rate-homeostatic rule, tested in the presence of strongly correlated assemblies or long-tailed weight distributions, can also generate a segregated structure with correlation-dependent selectivity (Vogels et al. 2011; Lagzi et al. 2021; Lagzi and Fairhall 2024). To reveal this effect, we consider a motif of one excitatory and two inhibitory neurons that both regulate the activity of the excitatory neuron through plastic synapses (Fig. S4, left). In line with the example in the main text, the inhibitory neurons are held at a constant rate *r*_inh_, while the excitatory firing depends on a fixed input and on the inhibitory feedback. The excitatory neuron projects to only one of the inhibitory neurons with a strong weight *w*_exc_, so that inhibitory neuron 1 is strongly correlated to it, but inhibitory neuron 2 is not. Calling *w*_1_ and *w*_2_ the inhibitory-to-excitatory weights, and considering them as plastic, we can write the weight dynamics as a simple generalization of Eq. S16:

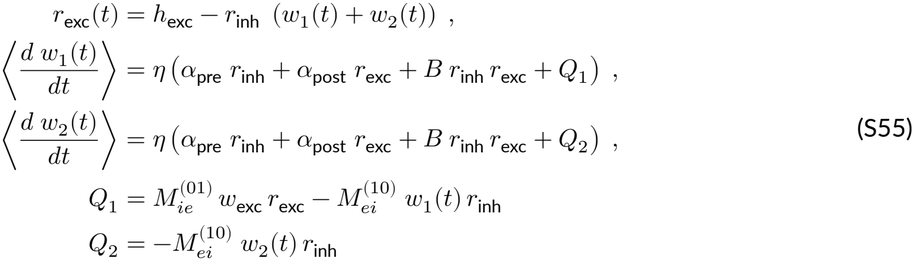

Here η is the global learning rate, and *B* is the area of the STDP kernel. The covariance terms *Q*_1_ and *Q*_2_ retain only first-order motif contributions, as in Eq. S31.

We consider two symmetric plasticity rules, either a rate-dominated rule, with *B*^(rate)^ = 1, or a covariance-dominated rule, with *B*^(cov)^ = 0. For both rules we set *α*_post_ = 0 and choose *α*_pre_ so that *r*_exc_(*t* → ∞) = *r*_target_. We then choose the learning rate η to give the same prescribed relaxation time *τ*_target_ for *r*_exc_(*t*) in both rules.

To solve Eq. S55, we first consider that the excitatory rate only depends on *w*_*s*_(*t*) := *w*_1_(*t*) + *w*_2_(*t*), resulting in a one-variable system:

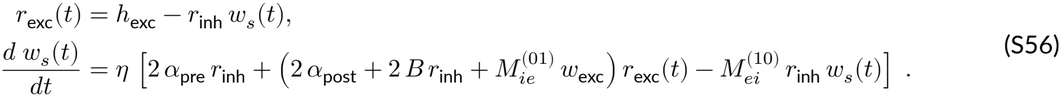

Equivalently, the excitatory rate obeys

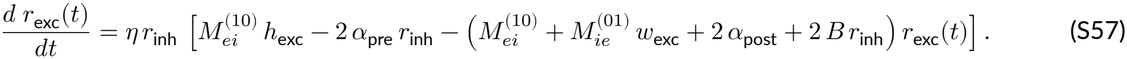

We can therefore express the fixed-point solution and the relaxation time in closed form:

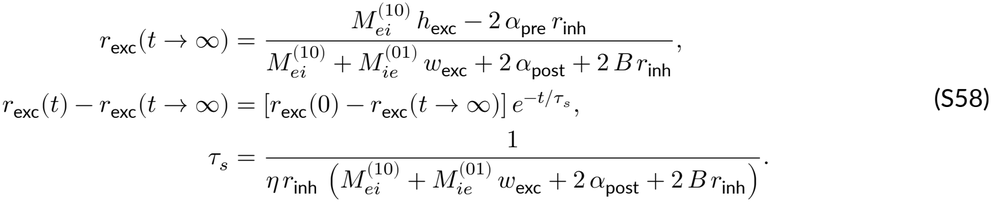

The remaining free parameters for the rate- and covariance-dominated rules are found by inverting Eq. S58 with the prescribed *τ*_target_ and *r*_target_:

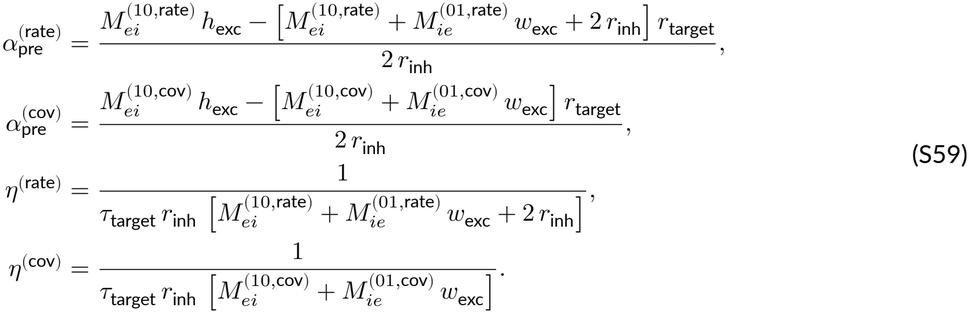

This choice sets *τ*_*s*_ = *τ*_target_ as the convergence timescale of the excitatory rate (and of the sum mode *w*_*s*_). The separation between the two inhibitory weights is captured instead by the difference mode *w*_*d*_(*t*) := *w*_1_(*t*) − *w*_2_(*t*), which obeys

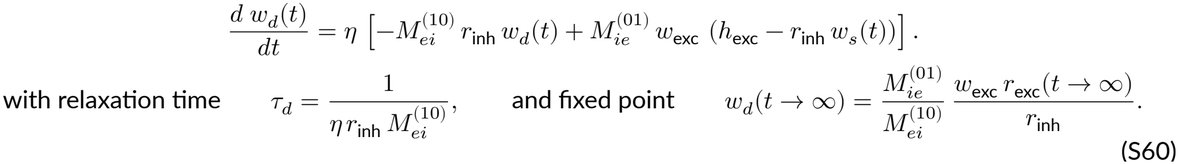

Note that all rules that satisfy 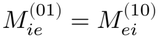 converge to the exact same fixed-point weights for a given *r*_target_. However, the convergence timescales of the two modes can differ substantially, as seen from their ratio:

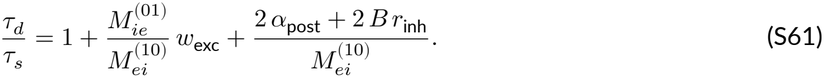

In the rate-dominated rule, where *B* = 1 and 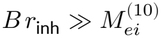, we obtain *τ* ≫ *τ* . By contrast, in the covariance-dominated rule, *B* = 0; for *α*_post_ = 0 and *w*_exc_ ∼ *O*(1), the two timescales remain of the same order (*τ*_*d*_ ∼ *τ*_*s*_).

Numerical simulations of the two rules confirm our analytical predictions (Fig. S4). In both cases, the excitatory rate approaches *r*_target_ on the prescribed timescale *τ*_target_ (Eq. S58), and the weights converge to the same steady-state solution (Eq. S60). However, in line with Eq. S61, under the rate-dominated rule the separation between *w*_1_ and *w*_2_ occurs on a much slower timescale than rate convergence (Fig. S4, green lines), whereas the covariance-dominated rule stabilizes the rate and separates the weights simultaneously (Fig. S4, purple lines).

## S3 Structured stabilization in spiking networks with intrinsically-generated fluctuating activity

Our main numerical results are based on conductance-based spiking RNNs, where excitatory neurons receive random Poisson inputs, with both a shared and an independent component. To further demonstrate the generality of our results, we consider a somewhat different but important class of spiking RNNs, where fluctuating activity is generated completely intrinsically and does not depend on external inputs (Van Vreeswijk and Sompolinsky 1998; Mastrogiuseppe and Ostojic 2017).

We adapted a current-based spiking RNN model that spontaneously generates irregular activity under constant input currents (Mastrogiuseppe and Ostojic 2017). The single-neuron dynamics is as follows:

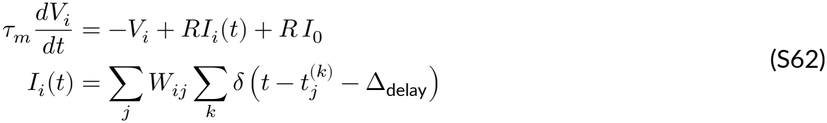

(see Table S1 for the full list of parameters). We selected initial conditions that resulted in a high-firing, nearly synchronous firing regime (Fig. S6A,B). In line with our previous results, we found that our choices of iSTDP rules lead to stable dynamics, reducing excitatory firing rates. Moreover, depending on the profile of correlation kernel, the network converged to either a prevalence of reciprocal E/I connections Fig. S6E, or lateral connectivity, expressed by unidirectional connections Fig. S6F.

These observations underscore the robustness of our proposed iSTDP mechanisms in shaping network connectivity beyond specific network architectures. The consistent emergence of either reciprocal or lateral connections, depending on the iSTDP rule, highlights a general principle by which local plasticity can shape E/I circuit structure.

**Table S1:**
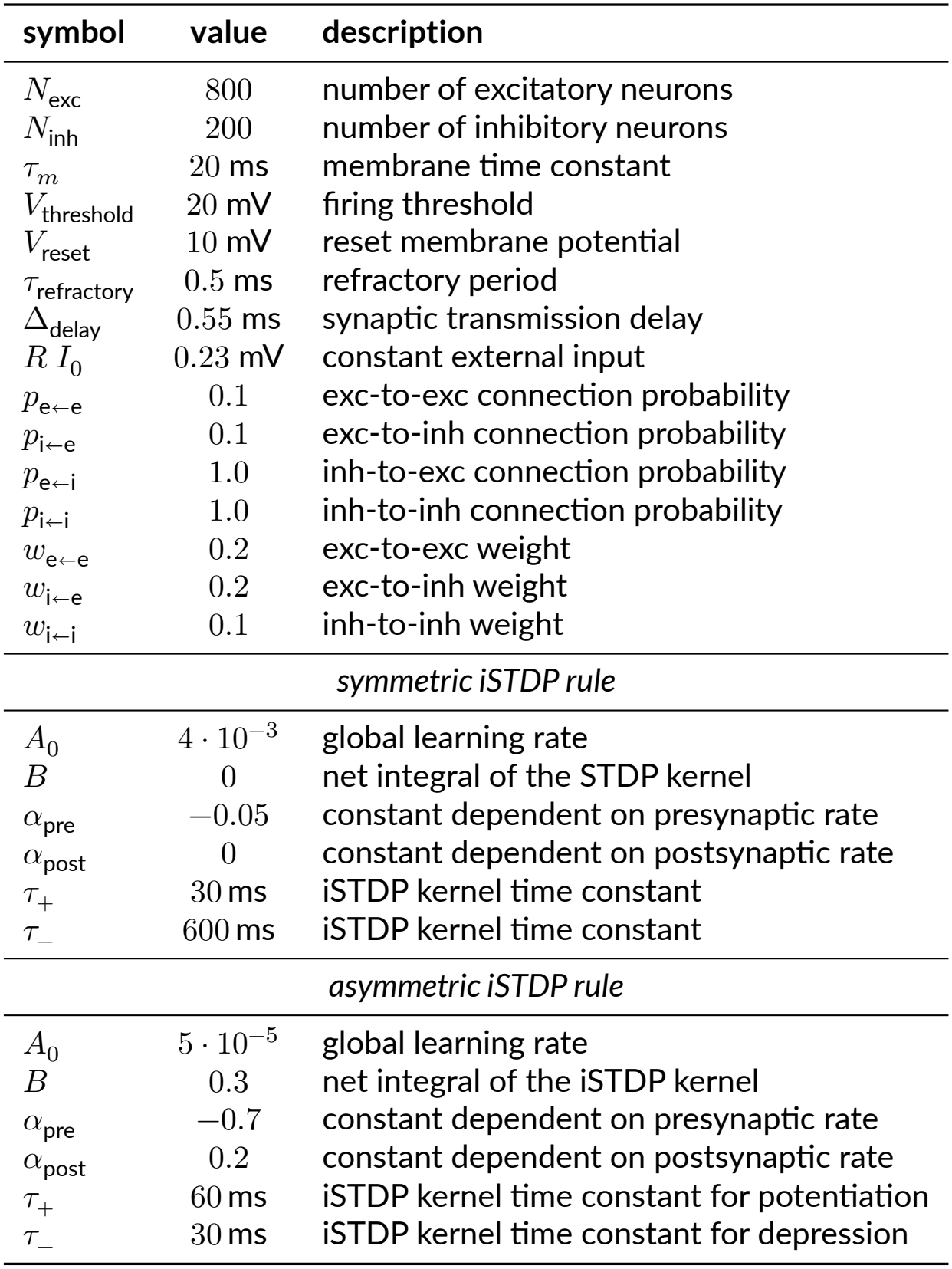
Numerical parameters for the spiking RNN with intrinsically-generated fluctuating activity (Eq. S62)

| symbol | value | description |
|---|---|---|
| N<sub>exc</sub> | 800 | number of excitatory neurons |
| N<sub>inh</sub> | 200 | number of inhibitory neurons |
| τ<sub>m</sub> | 20 ms | membrane time constant |
| V<sub>threshold</sub> | 20 mV | firing threshold |
| V<sub>reset</sub> | 10 mV | reset membrane potential |
| τ<sub>refractory</sub> | 0.5 ms | refractory period |
| Δ<sub>delay</sub> | 0.55 ms | synaptic transmission delay |
| R I<sub>0</sub> | 0.23 mV | constant external input |
| p<sub>e←e</sub> | 0.1 | exc-to-exc connection probability |
| p<sub>i←e</sub> | 0.1 | exc-to-inh connection probability |
| p<sub>e←i</sub> | 1.0 | inh-to-exc connection probability |
| p<sub>i←i</sub> | 1.0 | inh-to-inh connection probability |
| w<sub>e←e</sub> | 0.2 | exc-to-exc weight |
| w<sub>i←e</sub> | 0.2 | exc-to-inh weight |
| w<sub>i←i</sub> | 0.1 | inh-to-inh weight |
| *symmetric iSTDP rule* |  |  |
| A<sub>0</sub> | 4 · 10<sup>−3</sup> | global learning rate |
| B | 0 | net integral of the STDP kernel |
| α<sub>pre</sub> | −0.05 | constant dependent on presynaptic rate |
| α<sub>post</sub> | 0 | constant dependent on postsynaptic rate |
| τ<sub>+</sub> | 30 ms | iSTDP kernel time constant |
| τ<sub>−</sub> | 600 ms | iSTDP kernel time constant |
| *asymmetric iSTDP rule* |  |  |
| A<sub>0</sub> | 5 · 10<sup>−5</sup> | global learning rate |
| B | 0.3 | net integral of the iSTDP kernel |
| α<sub>pre</sub> | −0.7 | constant dependent on presynaptic rate |
| α<sub>post</sub> | 0.2 | constant dependent on postsynaptic rate |
| τ<sub>+</sub> | 60 ms | iSTDP kernel time constant for potentiation |
| τ<sub>−</sub> | 30 ms | iSTDP kernel time constant for depression |

## S4 Supporting Figures

**Figure S1:**
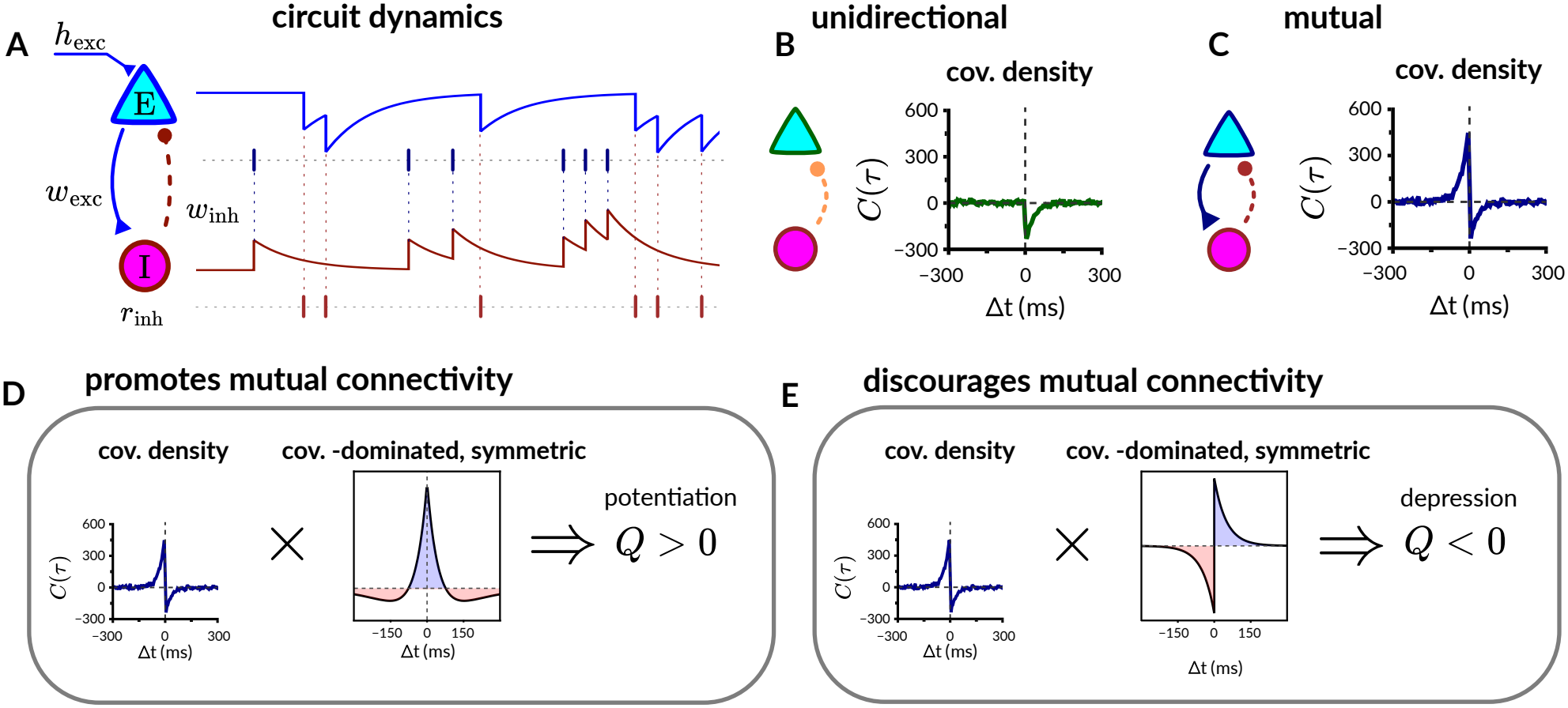
Schematics of the reduced circuit model and of interactions between covariance density and iSTDP kernel. **A**. Representation of the model used for Fig. 1 in main text. Two time-varying coupled Poisson units, one excitatory and one inhibitory, are connected by a non-plastic excitatory weight *w*_exc_ and a plastic inhibitory weight *w*_inh_. The instantaneous rates, here shown as traces, are computed as the convolution between each incoming spike and an exponential synaptic kernel that either increases (exc) or decreases (inh) the instantaneous firing probability. The input to the excitatory neuron *h*_exc_ is fixed, whereas the inhibitory input changes dynamically, keeping the inhibitory neuron at a fixed rate *r*_inh_. Only the inhibitory synapse (dashed line) is plastic. **B**. Unidirectional configuration. Here *w*_exc_ = 0 and the covariance density between the pre (inh) and post (exc) neuron can only be negative in the pre-post (Δ*t* > 0) section of the curve, due to the inhibitory interaction given by *w*_inh_. **C**. Mutual configuration, *w*_exc_ = 0.5. Here the covariance density is positive in the post-pre (Δ*t* < 0) section of the curve due to *w*_exc_, and negative for (Δ*t* > 0) for the effect of *w*_inh_. **D**. Effect of a symmetric, covariance-dominated iSTDP kernel on a circuit in the M configuration: the positive left branch of the covariance density interacts with the kernel, resulting in a positive *Q* factor in the weight update. (Eq. 1). **E**. Effect of an antisymmetric iSTDP kernel. Here the positive covariance density interacts with the negative branch of the kernel, resulting into synaptic depression. Hence, the *Q* term is negative, and mutual connections are discouraged.

**Figure S2:**
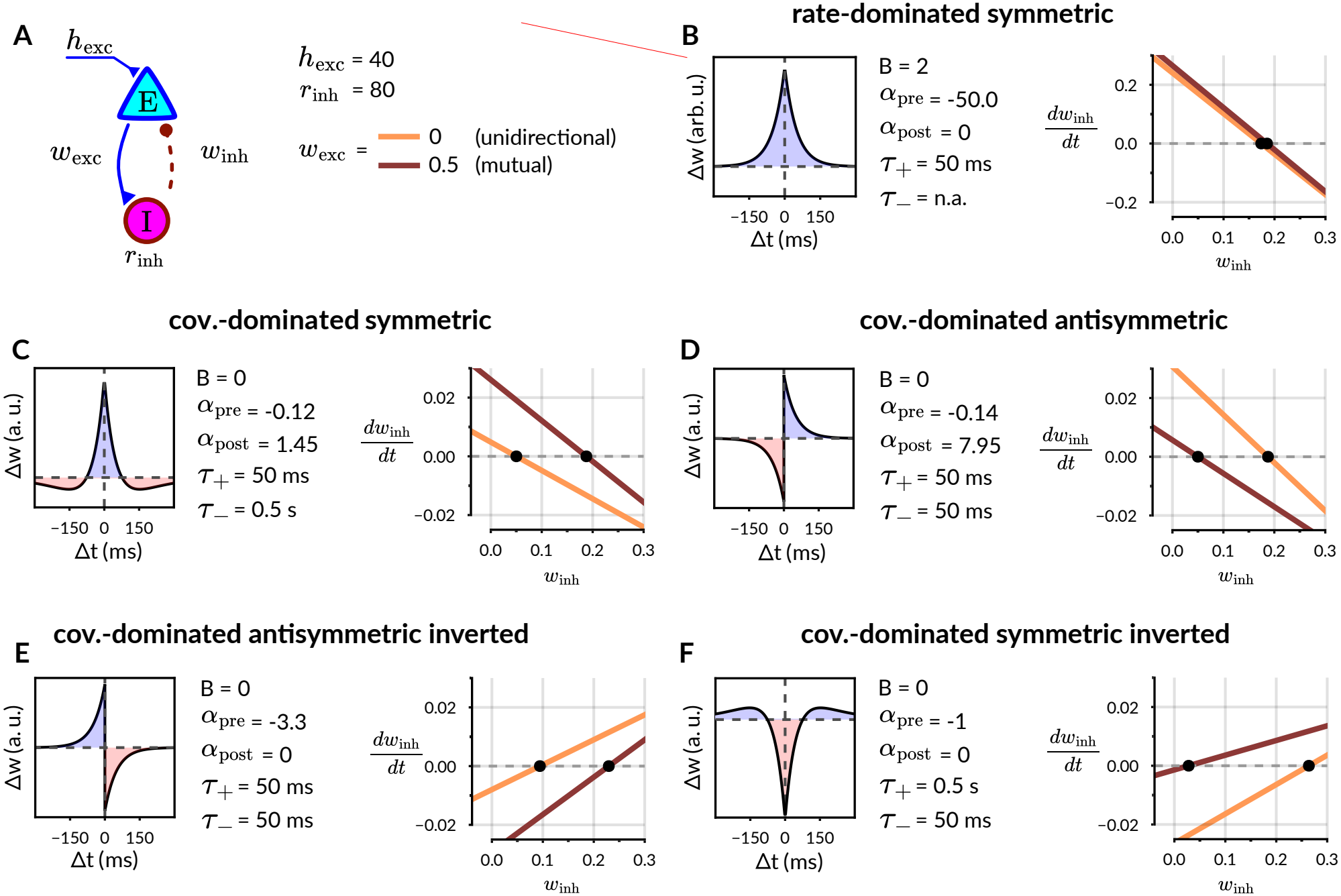
Analytic fixed point solutions for different iSTDP interaction kernels (complements Fig. 2 in main text). **A**. Representation of the model (see also Fig. 1A). **B-F**. iSTDP rules, represented by their kernel and their additional parameters (left panels) and study of the associated weight dynamics as a one-dimensional differential equation (right panels), black dots represent fixed points. **B** Rate-dominated iSTDP rule. **C-F**. covariance dominated rules presented in the same order as in Fig. 2A, main text. Note that the “saturating” configurations in **E** and **F** produce unstable fixed points (*dW* /*dt* has a positive slope at the zero-crossing), so weights diverge to 0 or *w*_max_.

**Figure S3:**
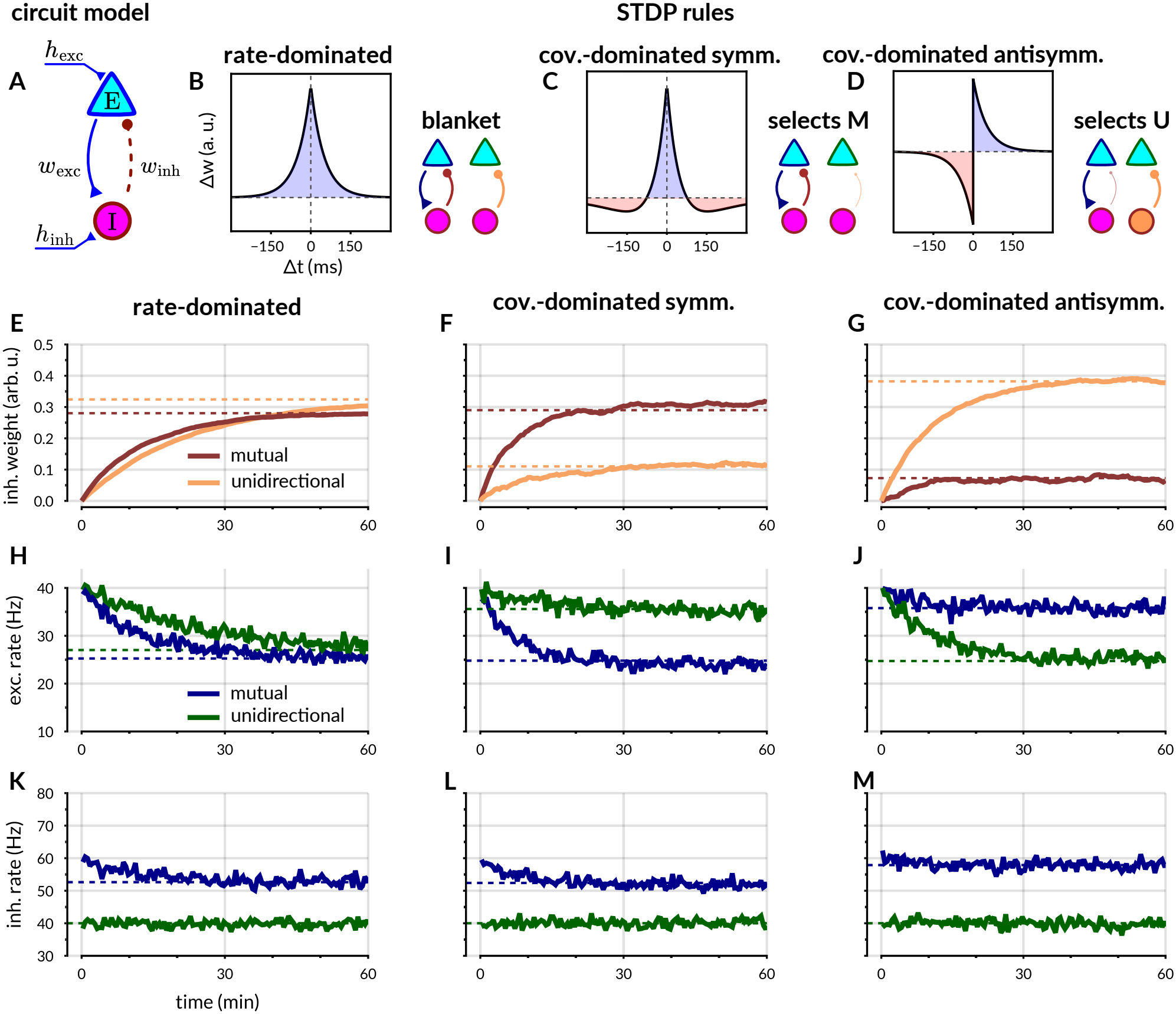
Effects of plasticity for E/I motifs in a reduced model with unconstrained *r*_inh_. See Section S1.3 for model description and details on the analytic calculation. **A**. Model schematics. Unlike the model in Fig. 1A, the inhibitory rate *r*_inh_ is unconstrained and determined by the net inhibitory input *h*_inh_. **B-J**. Corresponding to the respective panels in Fig. 1, for the model with unconstrained *r*_inh_. **K-L**. Value of *r*_inh_ during the simulation. Note that in the U configuration (dark green) *r*_inh_ is stationary and corresponds to *h*_inh_.

**Figure S4:**
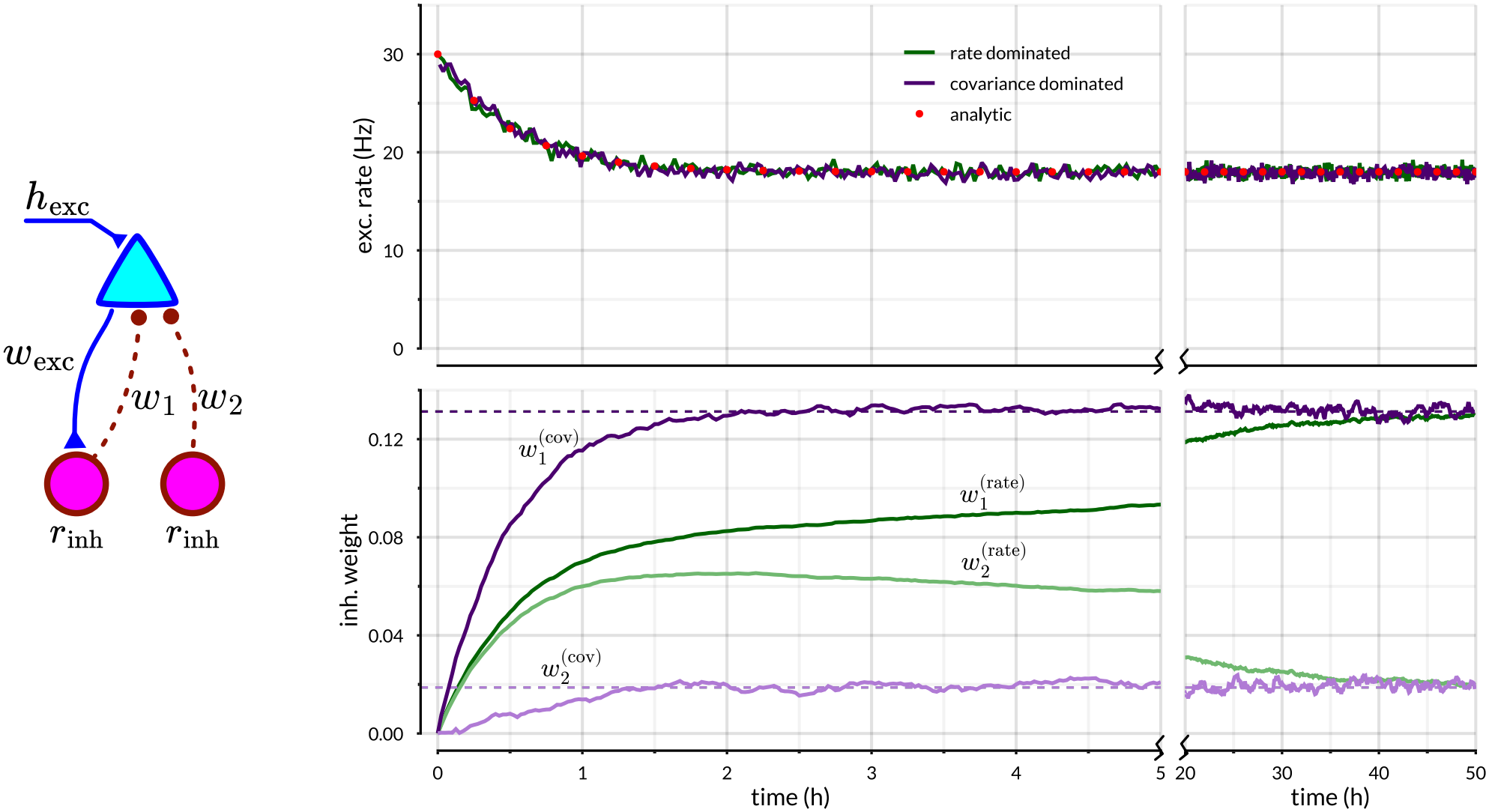
Rate-dominated and covariance-dominated iSTDP in a three-neuron motif (see Section S2). **Left**. One excitatory neuron receives a fixed input *h*_exc_ and inhibition through two plastic weights, *w*_1_ and *w*_2_, from neurons held at rate *r*_inh_ by an adaptive input. The excitatory neuron projects to the first inhibitory neuron with a fixed weight *w*_exc_. **Right, top**. Excitatory rate under the rate-dominated (green) and covariance-dominated (purple) rules, calibrated to the same target rate *r*_target_ and relaxation time *τ*_target_. Red dots show the analytic prediction. **Right, bottom**. Evolution of *w*_1_ (dark) and *w*_2_ (light) for the rate-dominated (green) and covariance-dominated (purple) rules. Dashed lines show the analytic fixed-point weights. The two rules approach the same final weights, but the covariance-dominated rule separates the inhibitory weights and establishes a mutually connected E/I motif on a shorter timescale.

**Figure S5:**
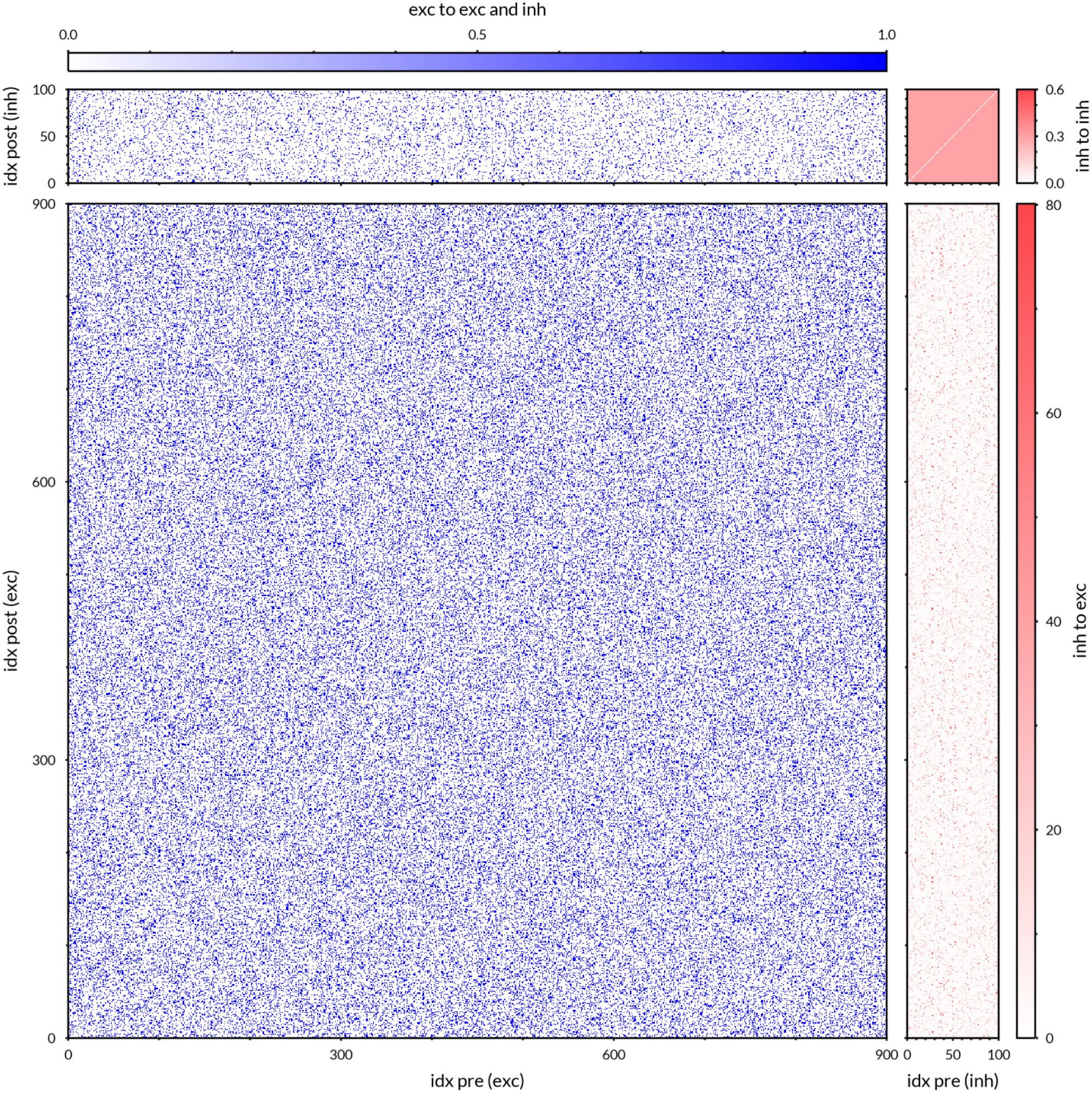
Full weight matrix for the random sparse RNN simulation with symmetric iSTDP (see Main Text, Fig. 3B-F).

**Figure S6:**
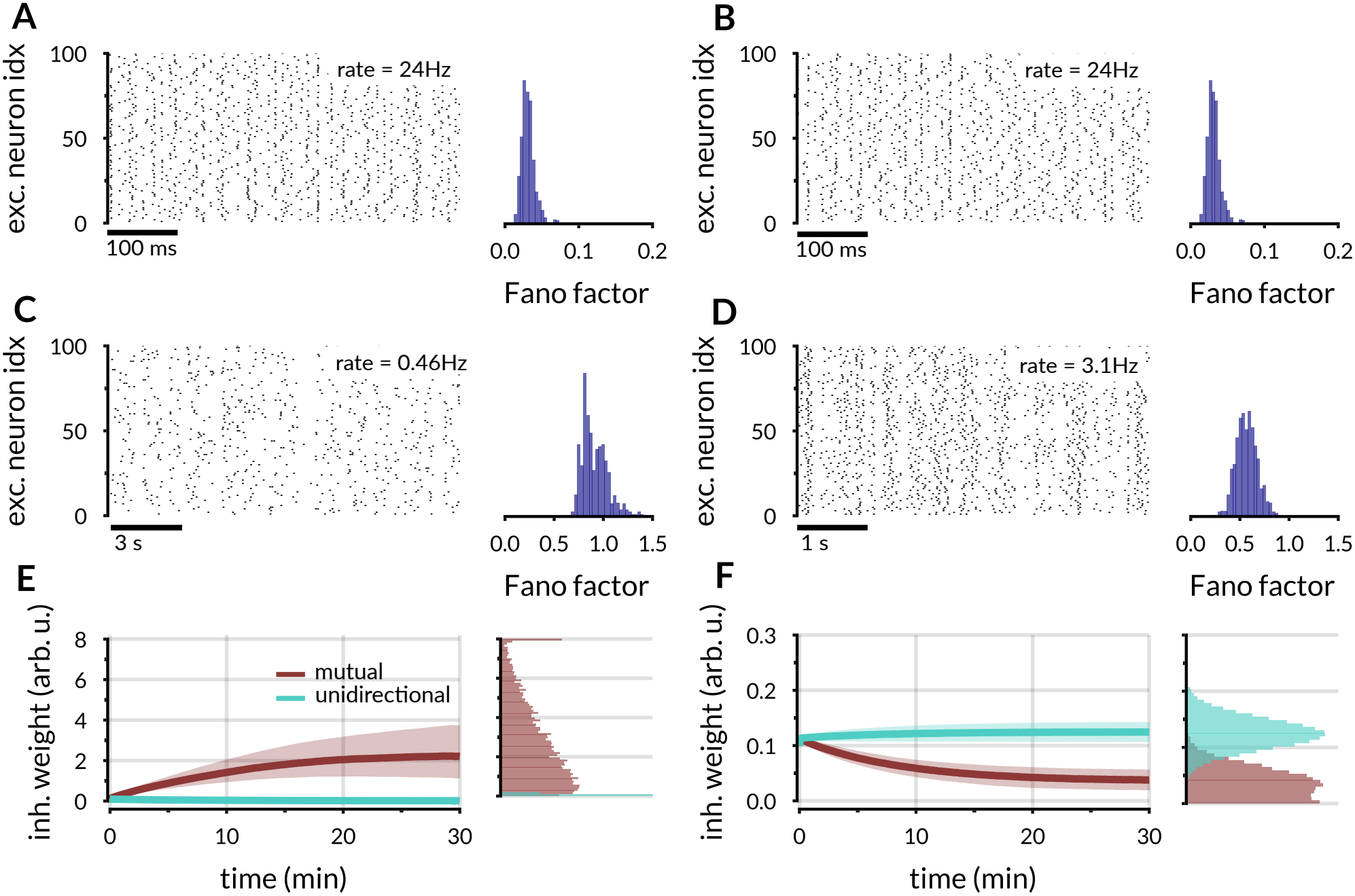
Randomly connected spiking RNN with intrinsically-generated fluctuating asynchronous irregular activity. See Table S1 for numerical parameters. **A-B**. Spiking activity of a subset 100 excitatory neurons (left) and distribution of Fano factors of excitatory neurons (right) before plasticity. **C-D**. Spiking activity and Fano factor distribution after weight convergence, using either a symmetric (**C**) or an antisymmetric (**D**) iSTDP kernel. **E-F**. Change of mutual and unidirectional inhibitory-to-excitatory weights during learning in the circuit. Equivalent to Fig. 3C and H in Main Text. Thick lines show the median and shaded areas span the 25th to 75th percentiles of the synaptic weight distributions.

**Figure S7:**
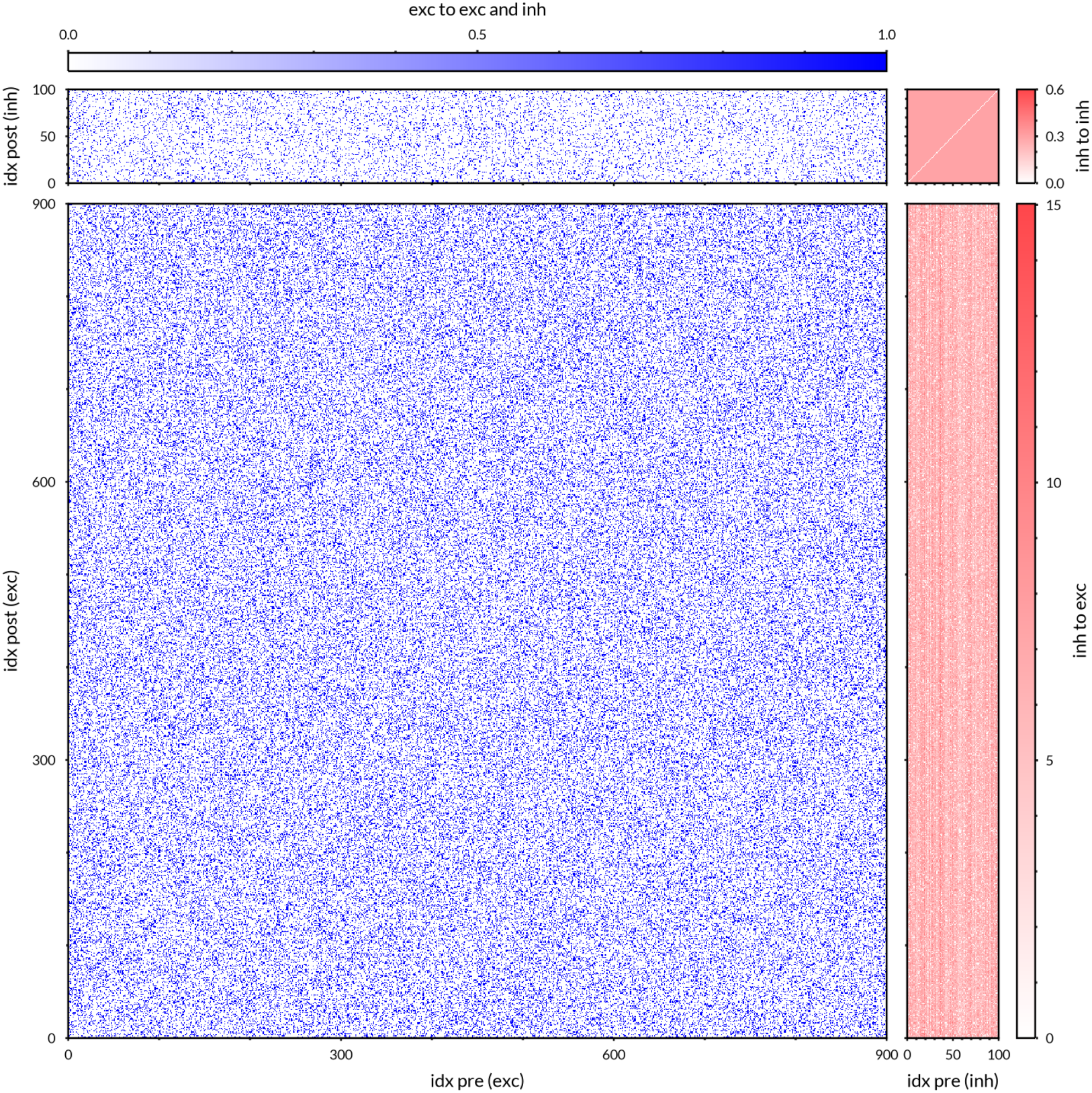
Full weight matrix for the random sparse RNN simulation with antisymmetric iSTDP (see Main Text, Fig. 3G-K).

**Figure S8:**
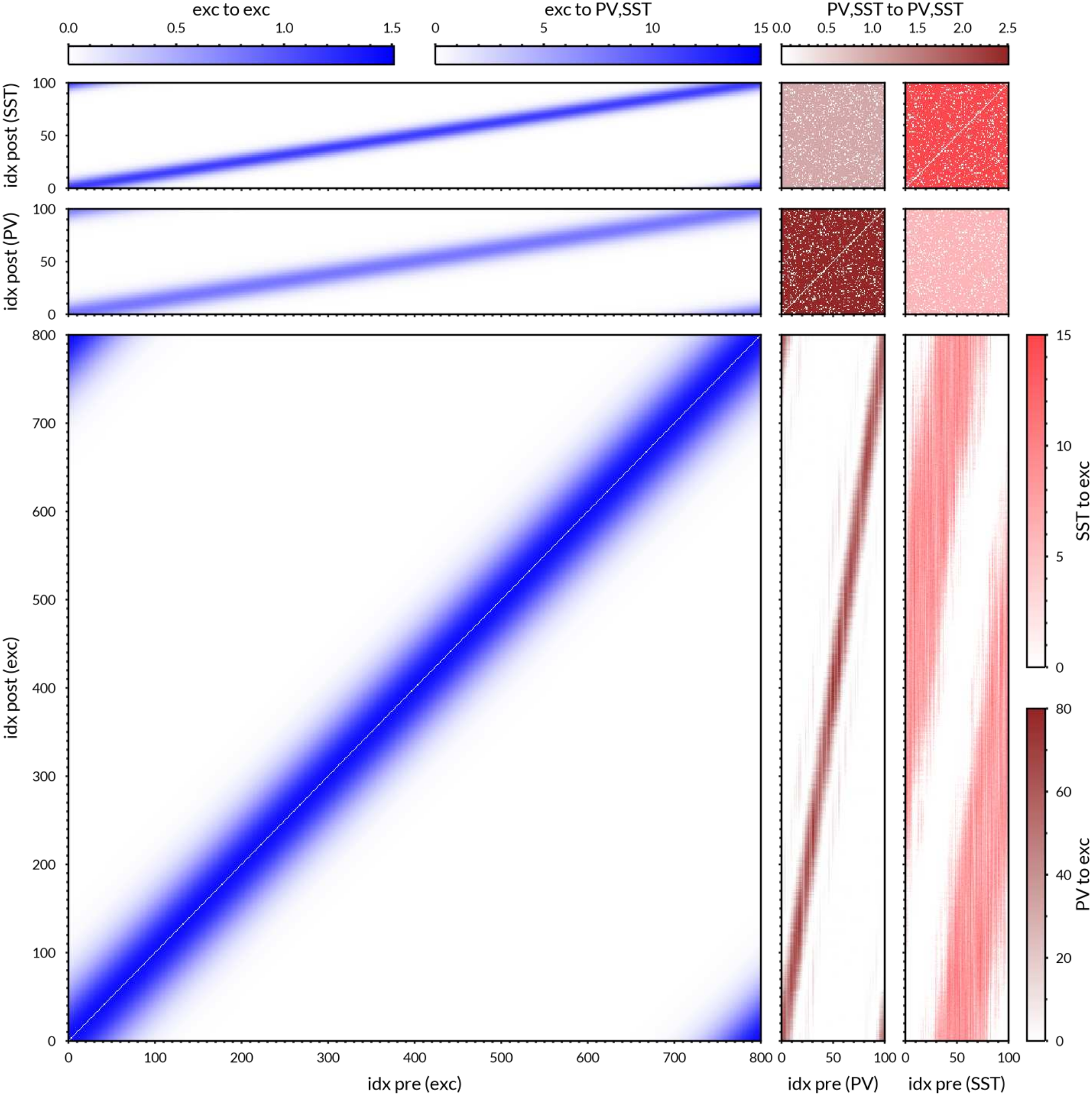
Full weight matrix for the recurrent network with 1D ring connectivity structure after weight convergence with two inhibitory populations.

**Figure S9:**
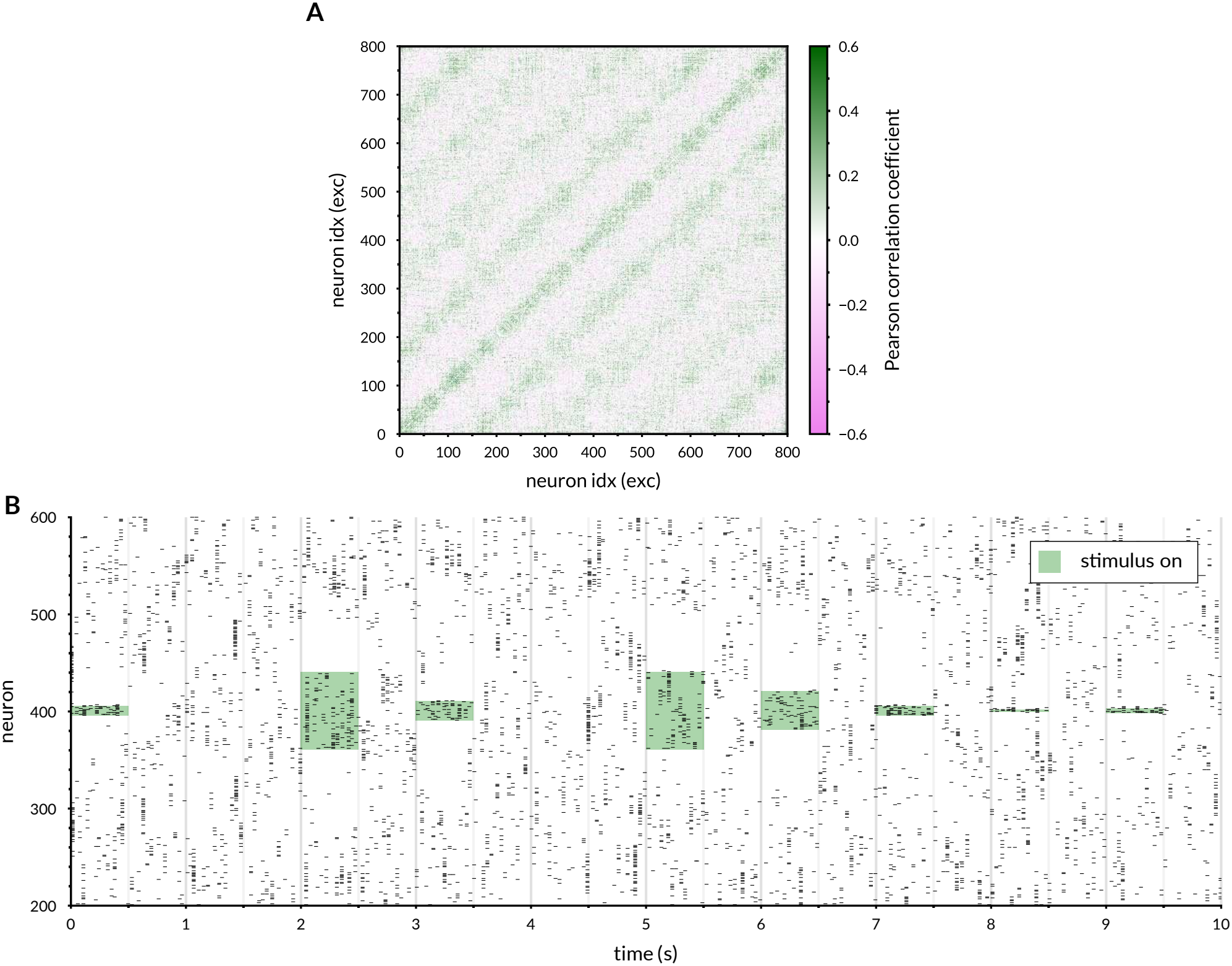
**A**. Full matrix of Pearson correlations between excitatory neurons in the one-dimensional ring model during spontaneous activity, after convergence of plastic weights. **B** Spike raster showing the response of the 1D ring model to localized stimuli (see Fig. 4 in main text).

**Figure S10:**
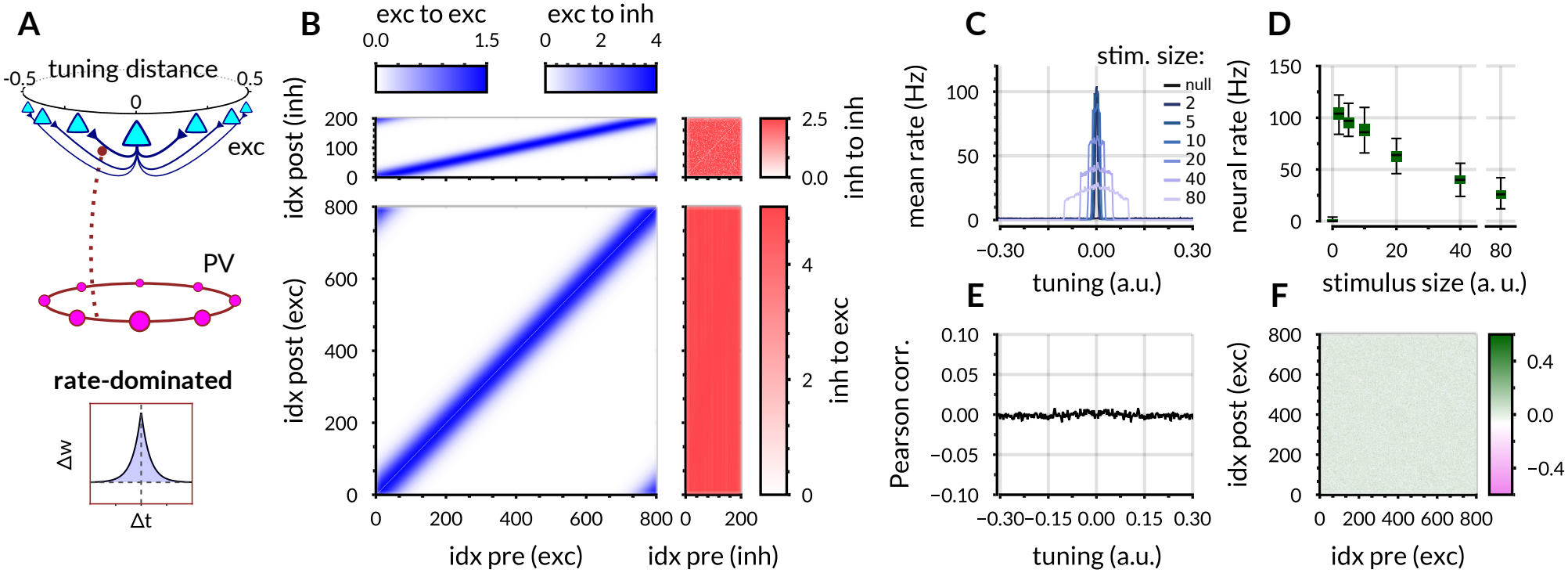
Spontaneous and evoked activity in a 1D ring structure with rate-dominated iSTDP. **A**. Model schematics, compare with Fig. 4A. The excitatory component is identical to the previous case, but the inhibitory population follows a single, rate-dominated iSTDP. **B**. Full weight matrix after training, compare with Fig. 4B. **C**. Mean responses of the excitatory population to stimuli of different sizes, averaged over 100 trials (compare with Fig. 5A). **D**. Response of the center neuron to stimuli of different sizes (compare with Fig. 5B). **E**. Pearson correlation during spontaneous activity between an excitatory neuron and its neighbors, ordered by tuning distance and averaged over the entire population (compare with Fig. 4F). **F**. Pearson correlation of spontaneous activity for every pair of excitatory neurons (compare with Fig. S9A). Note that over much longer simulation times, slow weight drift may still lead to the formation of structured connectivity (see Section S2 and Fig. S4).

**Figure S11:**
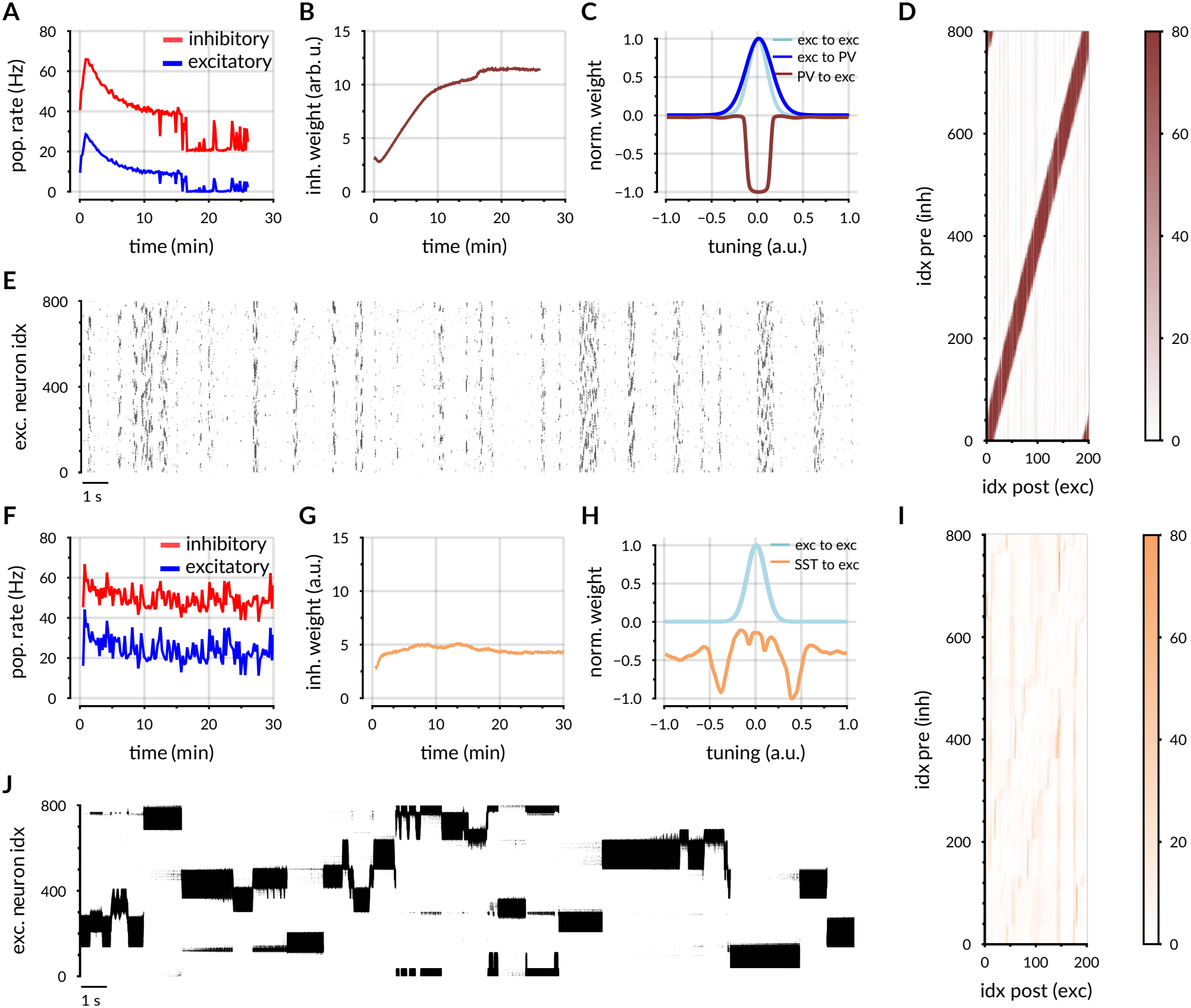
One-dimensional ring network with only one inhibitory population (either PV or SST). **A**-**E**. The RNN has 800 excitatory neurons, 200 “PV” neurons and no “SST” neurons. The network evolves under a PV-like covariance-dominated iSTDP rule with *B* = 0, *α*_pre_ = −1.0, *α*_post_ = 1.0. **A**. Evolution of population mean firing rates over time. **B**. Evolution of inh-to-exc synaptic weights over time. **C**. Profile of the average outgoing weights as a function of tuning distance. For convenience, the profiles are normalized so that the maximum is either 1 or −1. **D**. Inhibitory to excitatory weight matrix after weight convergence. **E**. Raster plot of excitatory neurons. **F**-**J**. Same but for a RNN with 800 excitatory neurons, 200 “SST” neurons and no “PV” neurons. The network evolves under a SST-like covariance-dominated iSTDP rule with *B* = 0, *α*_pre_ = −1.0, *α*_post_ = 1.5.

**Figure S12:**
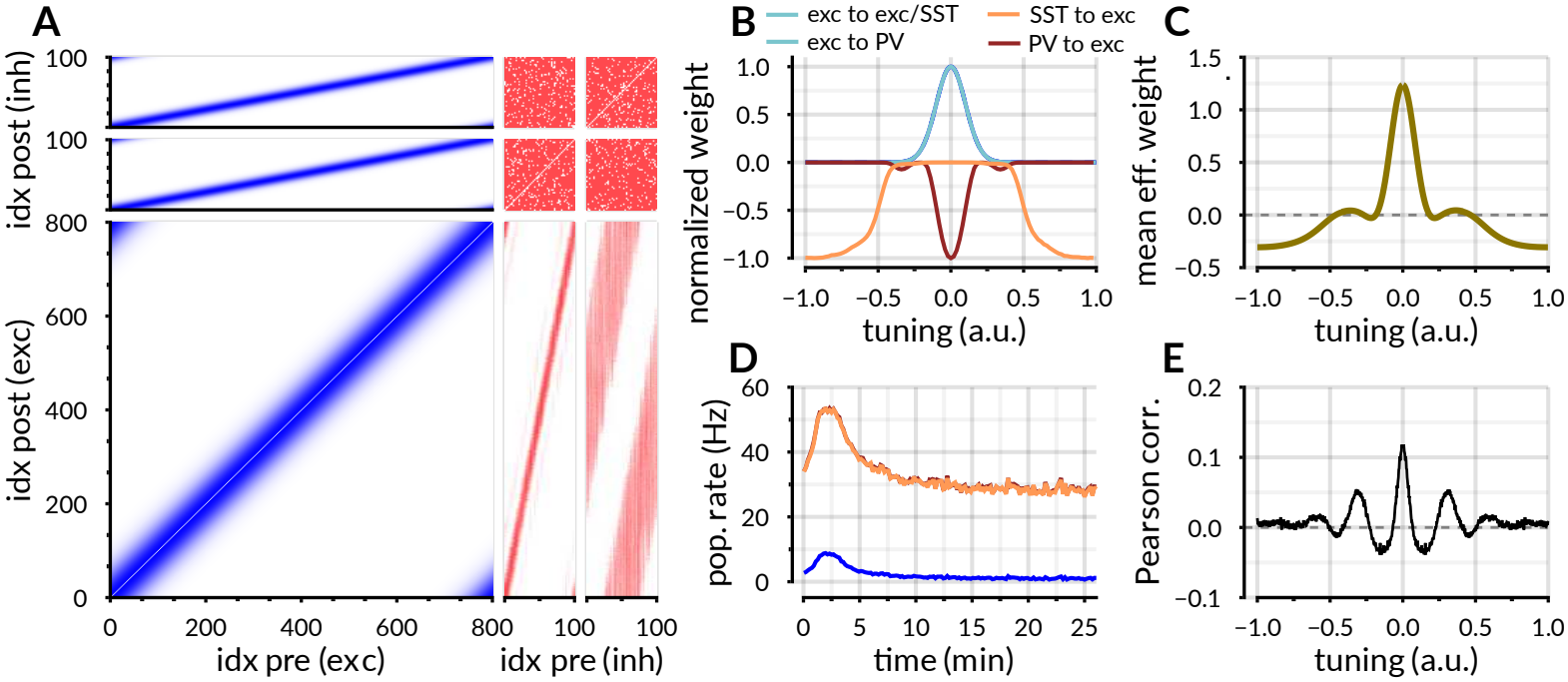
One-dimensional ring network in which all outgoing excitatory connections have the same spatial range. The spatial range of excitatory-to-PV connections is the only parameter that differs from the model in Fig. 4. Panels **A**-**E** correspond to panels **B**-**F** in Fig. 4. The main difference observed is a faster decay of long-range correlations (compare panel **E** with Fig. 4F)

**Figure S13:**
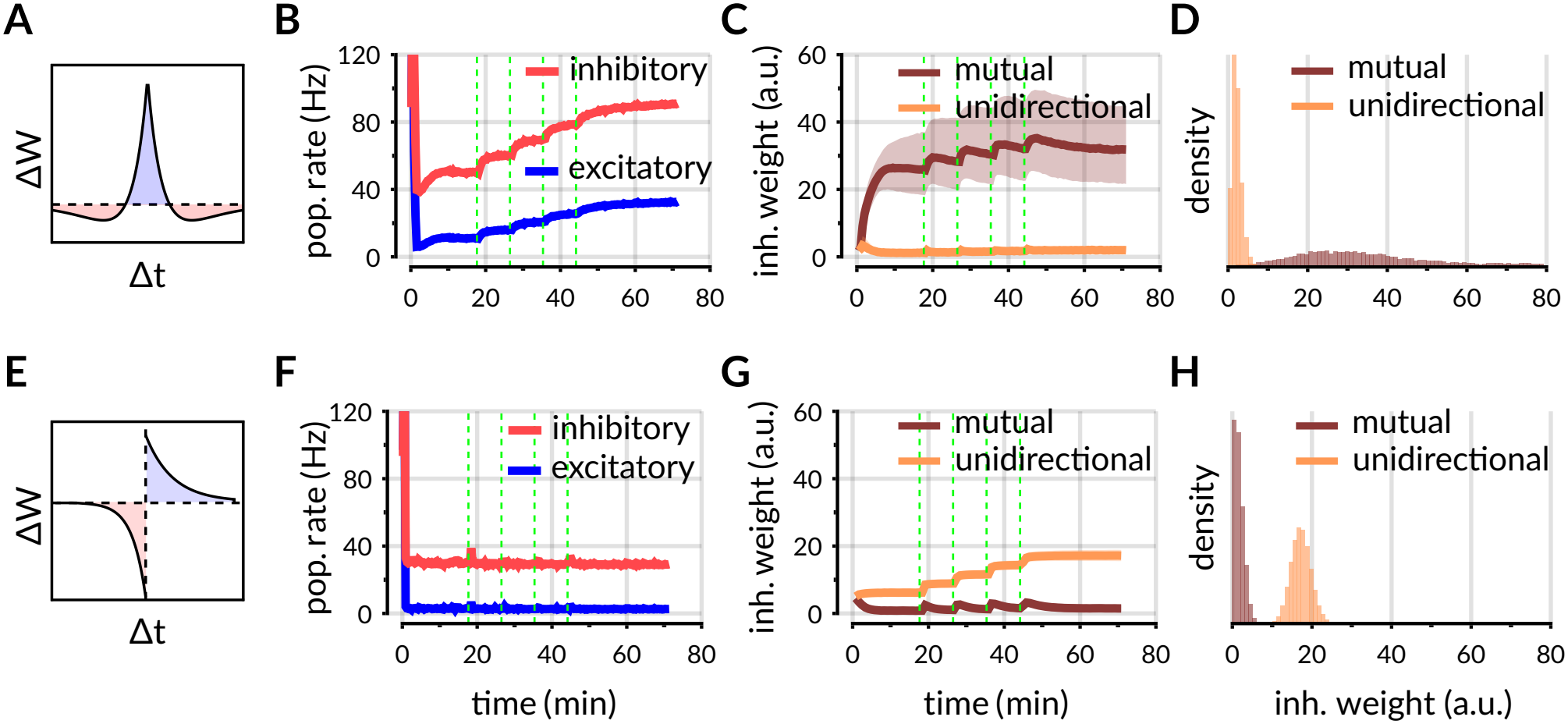
Effects of plasticity for different levels of mean inhibitory input. **A**. Symmetric kernel. **B**. Evolution of population mean firing rates over time. The green vertical lines represent sequential increases of the mean input by a factor of 1.5, 2.0, 2.5, 3. **C**. Evolution of inh-to-exc synaptic weights over time. **D**. Final distribution of weights, split into a “mutual” (brown) and a “unidirectional” (light orange) group. **E-H**. Same as above but for asymmetric iSTDP. In panels **C** and **G**, thick lines show the median and shaded areas span the 25th to 75th percentiles of the synaptic weight distributions.

